# Mesoscopic analysis of GABAergic marker expression in acetylcholine neurons in the whole mouse brain

**DOI:** 10.1101/2025.09.10.675394

**Authors:** R. Oliver Goral, Snehashis Roy, Caroll A. Co, Robert N. Wine, Patricia W. Lamb, Peyton M. Turner, Sandra J. McBride, Ted B. Usdin, Jerrel L. Yakel

**Affiliations:** Neurobiology Laboratory, National Institute of Environmental Health Sciences, National Institutes of Health, Department of Health and Human Services, Research Triangle Park, NC, USA; Center on Compulsive Behaviors, National Institutes of Health, Bethesda, MD, USA; Systems Neuroscience Imaging Resource, National Institute of Mental Health, National Institutes of Health, Department of Health and Human Services, Bethesda, MD, USA; DLH, LLC, Bethesda, MD, United States; Fluorescence Microscopy Imaging Center, National Institute of Environmental Health Sciences, National Institutes of Health, Department of Health and Human Services, Research Triangle Park, NC, USA

## Abstract

In the central nervous system, acetylcholine (ACh) neurons coordinate neural network activity required for higher brain functions, such as attention, learning, and memory, as well as locomotion. Disturbances in cholinergic signaling have been described in many diseases of the developing and mature brain. Interestingly, ACh neurons can co-transmit GABA to support essential roles in brain function. However, the contributions of ACh/GABA co-transmission to brain function remain unclear. This underscores the need to better understand the heterogeneity of ACh neurons, particularly the sub-population of ACh neurons co-expressing GABAergic markers. We used various combinations of transgenic mouse lines to systematically label ACh neuron populations positive for different GABAergic markers in the brain. We developed a workflow combining tissue clearing, light-sheet fluorescence microscopy, and machine learning to image entire mouse brain hemispheres followed by quantification of ACh neurons throughout the brain. With this approach, we assessed the role of GABA co-transmission in ACh neuron function and quantified ACh and ACh/GABA neuron sub-populations in the brain. Our results suggest that GABA co-transmission from ACh neurons is not required to maintain the regular ACh neuron count in the brain. Furthermore, we report that a large subset of ACh neurons can potentially synthesize GABA by co-expressing the marker Gad2. However, most of these ACh neurons do not express vGAT, which would enable these neurons to release GABA. Taken together, we conclude that GABA co-transmission likely occurs only from a small population of ACh neurons restricted to few brain nuclei.

**GRAPHICAL ABSTRACT:** 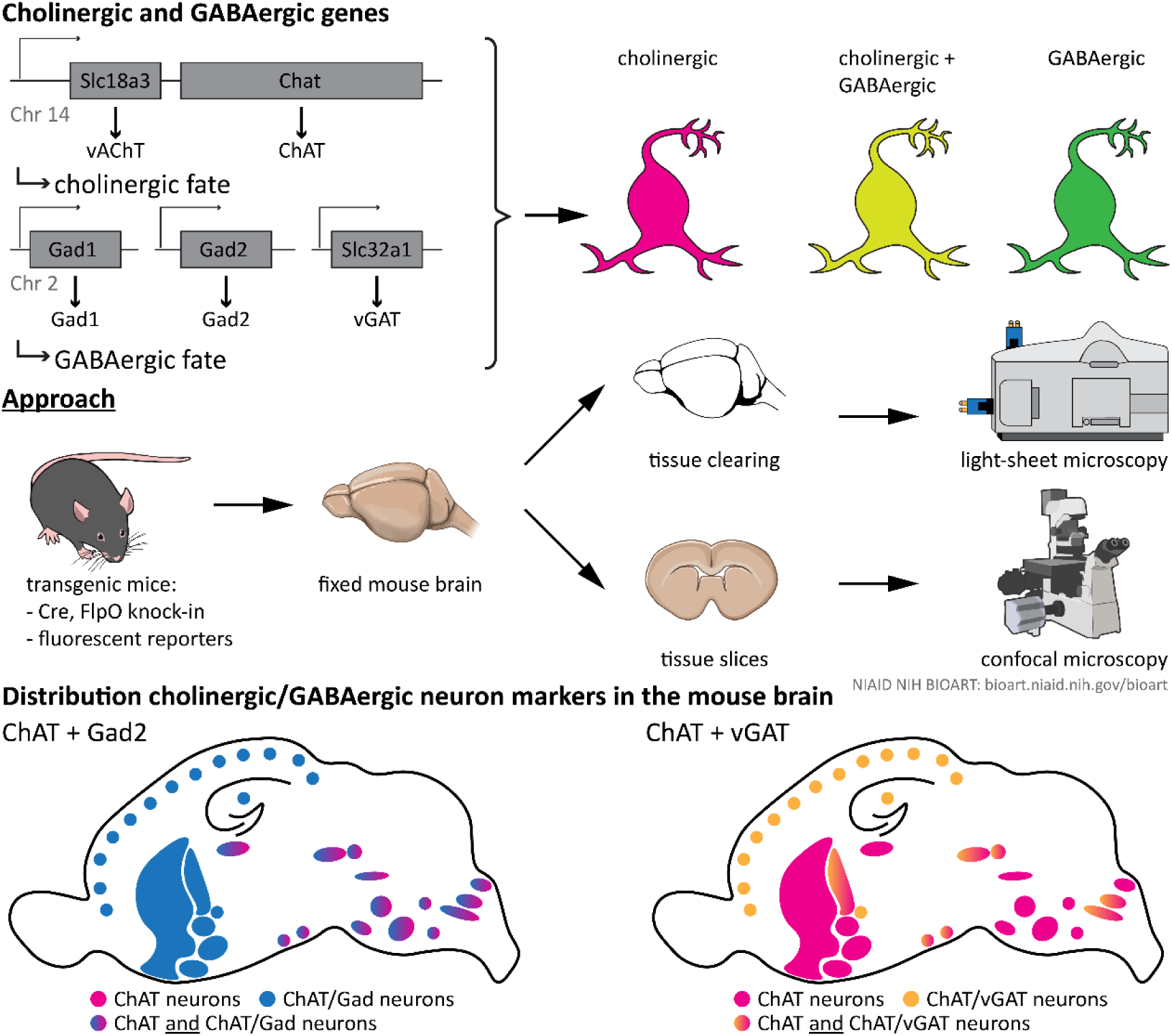

## Introduction

In the central nervous system, acetylcholine (ACh) neurons are essential for higher cognitive functions, such as attention, learning, and memory, as well as pattern generation in the spinal cord for locomotion (Ballinger et al. (2016), Mille et al. (2021)). ACh neurons or their projections can be found in every part of the brain (Li et al. (2018), Gamage et al. (2023), Goral et al. (2024)). The cholinergic system does not only moderate neuron firing and synaptic strength, but also neuron morphology, as well as the release of neuromodulators, such as dopamine (DA) or norepinephrine from axonal varicosities (Lozada et al. (2012), Morley and Mervis (2013), Liu et al. (2022), Cools and Arnsten (2022), Steinecke (2022)). With ACh neurons forming synapses on many neurons, they can provide accurately timed modulation of neuronal firing, which is believed to regulate oscillatory network activity in various brain regions required for the computation of higher brain functions (Marrosu (1995), Hasselmo and McGaughy (2004), Gu and Yakel (2011), Unal et al. (2015), Gu et al. (2017), Gu et al. (2020), Gu et al. (2024)).

Co-release or co-transmission of multiple signaling molecules at the same synapse has been previously described as a distinctive mechanism that increases the complexity of synaptic computation (J L Ellis (1989), Wojcik et al. (2006), Upmanyu et al. (2022)). Many ACh neurons originate from the same proliferation zones and cell lineages, that generate GABAergic neuron populations (Ahmed et al. (2019), Ananth et al. (2023)). It is therefore not surprising that subsets of ACh neurons in forebrain and midbrain co-express GABAergic markers and co-transmit GABA at their synapses (Saunders et al. (2015a), Saunders et al. (2015b), Estakhr et al. (2017), Takacs et al. (2018), Lozovaya et al. (2018), Obermayer et al. (2019), Granger et al. (2020), Hunt et al. (2022), Le Gratiet et al. (2022), Granger et al. (2023), Lozovaya et al. (2023), Lozovaya et al. (2024)). GABA co-transmission from ACh neurons has been implied in network functions in hippocampus, striatum, and medial prefrontal cortex (mPFC), but the molecular mechanisms remain unknown (Takacs et al. (2018), Lozovaya et al. (2018), Obermayer et al. (2019), Granger et al. (2020)). Recent data suggests that the loss of GABA co-transmission from ACh neurons leads to impairments in brain functions, such as social, spatial, and fear memory as well as altered habit learning without affecting the overall health of the animals or their performance during more basic behaviors (Goral et al. (2022)). Although co-expression of GABAergic markers in ACh neurons peaks in the developing mouse brain during the second postnatal week and declines in adolescence, it remains unclear whether GABA co-transmission from ACh neurons is essential for brain development or instead serves acute modulatory functions in the brain (Granger et al. (2023), Lozovaya et al. (2023), Lozovaya et al. (2024)). Furthermore, the functional significance of GABA co-transmission in ACh neurons and the distribution of ACh/GABA neurons across the brain are still not well understood.

Therefore, we set out to quantify GABAergic and non-GABAergic ACh neuron sub-populations at the whole-brain level. For this purpose, we genetically labelled ACh neurons with the fluorescent marker tdTomato (tdT). These brains underwent tissue clearing and were imaged with a light-sheet fluorescence microscope. Brain tissue clearing has proven to be a powerful method for bypassing labor-intensive sectioning and post-hoc reconstruction in large volume imaging (Erturk et al. (2012), Chung et al. (2013), Renier et al. (2014), Renier et al. (2016), Richardson et al. (2021), Kosmidis et al. (2021)). We subsequently applied machine learning to segment and map ACh neurons across the whole brain. Here, we report that loss of GABA co-transmission from ACh neurons through the genetic ablation of the vesicular GABA transporter (vGAT) does not negatively affect ACh neuron numbers in the brain. Furthermore, we discovered that the fraction of GABAergic ACh neurons depends on the brain region and the GABAergic marker. Interestingly, we found the Gad2 co-positive ACh neuron sub-population is highly abundant throughout the forebrain, but less so in the mid-or hindbrain ACh nuclei. In contrast, the vGAT co-positive ACh neuron sub-population is restricted to few cholinergic nuclei indicating that expression of Gad2 and vGAT are regulated separately. Furthermore, we show that based on the combined expression pattern of these two GABAergic markers, GABA co-transmission can only occur from few cholinergic nuclei in the mouse brain.

## Material and Methods

### Experimental Design

No sample size pre-determination was conducted. At experiment onset, the goal was set to collect a total number of 8-10 cleared brain hemisphere datasets of sufficient quality per condition. For histology experiments, samples from at least three different animals were used for each condition. In addition, staining results were confirmed in several slices as well as by comparison with control slices (both unstained and secondary antibody only). Box whisker plots were constructed using Tukey’s method with outliers identified as data points exceeding 1.5x the interquartile range.

### Animals

All animal procedures were approved and performed in compliance with the NIEHS/NIH Humane Care and Use of Animals Protocols. All transgenic animal lines were purchased from The Jackson Laboratories and subsequently maintained and bred in-house. Animals were group housed (≤5 per cage) in a regular 12 h light/dark cycle under constant temperature control. Food and water were supplied *ad libitum*. We labelled ACh neurons by crossing homozygous ChAT-IRES-Cre (Jax# 006410, RRID:IMSR_JAX:006410, Rossi et al. (2011)) with homozygous Ai14 reporter mice (Jax# 007914, RRID:IMSR_JAX:007914, Madisen et al. (2010)) on a C57BL/6J background. To assess whether loss of GABA co-transmission from cholinergic neurons reduced ACh neuron count, we crossed vGAT-flox mice (Jax# 012897, RRID:IMSR_JAX: 012897), with ChAT-IRES-Cre mice and Ai14 mice (Tong et al. (2008)). Breeding was set up to obtain litters with mice either negative, homozygous, or heterozygous for vGAT-flox alleles and heterozygous for both Cre and tdT. Mice negative for floxed alleles were considered “vGAT WT”, while mice with homozygous floxed alleles were considered “vGAT CKO”. For comparison of tissue-specific recombinase expression, ChAT-FlpO mice were crossed with Ai65F FlpO reporter mice (Jax# 032864, RRID:IMSR_JAX:032864, Daigle et al. (2018)).

To further separate ACh neuron populations, ChAT-IRES-FlpO mice (Jax# 036281, RRID:IMSR_JAX:036281) were crossed with Gad2-IRES-Cre mice (Jax# 010802, RRID:IMSR_JAX:010802) and Ai65D Cre/FlpO reporter mice (Jax# 021875, RRID:IMSR_JAX:021875, Allaway et al. (2020), Taniguchi et al. (2011), Madisen et al. (2015)) on a C57BL/6J background. To test out another genetic separation strategy, we crossed vGAT-IRES2-FlpO-D (Jax # 031331, RRID:IMSR_JAX:031331) with ChAT-IRES-Cre and the Ai65D Cre/FlpO reporter (Daigle et al. (2018)). All animals used for experiments were heterozygous for all three transgenes, respectively. See Suppl. Fig. 1 for an overview of all used breeding schemes.

### Histology

We collected mouse brains between postnatal day (P) 55 and 65. Mice were deeply anesthetized with sodium pentobarbital (100 mg/kg) and transcardially perfused with 0.1 M PBS and 0.1 M PBS + 4% PFA and postfixed in 0.1 M PBS + 2% PFA at 4°C overnight. Afterwards, the brains were freeze-protected in 0.1 M PBS with 30% sucrose and frozen in Tissue-Tek O. C. T. Compound (Sakura Finetek, Torrance, CA, USA). For immunostaining, 50 µm slices were cut with a CM3050 cryostat (Leica, Deer Park, IL, USA) and stored in Tris-buffered saline (TBS) + 50% glycerol at -20°C until use. Slices were rinsed 3x in TBS. After washing, slices were blocked in TBS + 0.3% Triton X-100 (Thermo Fisher, Hillsboro, OR, USA) supplemented with 10% normal donkey serum (NDS, Abcam, Cambridge, MA, USA) or normal goat serum (NGS, Abcam) at RT for 2 hours. Afterwards, slices were incubated with primary antibodies diluted in TBS with 0.3% Triton X-100 and 5% NDS/NGS at 4°C overnight. After 3x washing in TBS, slices were incubated with secondary antibodies diluted in TBS with 0.3% Triton X-100 and 5% NDS/NGS at 4°C overnight. Slices were washed 3x in TBS before they were mounted on Superfrost Plus microscope slides (Thermo Fisher), air dried and protected with ProLong™ Glass Antifade Mountant with NucBlue™ (Hoechst 33342, Thermo Fisher) and thickness 1.0 microscope cover glasses.

### Fluorescence Confocal Microscopy

Fluorescent images were collected with an inverted LSM 980 laser scanning confocal microscope (Zeiss, Thornwood, NY, USA) controlled by the software Zen blue (RRID:SCR_013672, Zeiss). Fluorophores were excited with diode lasers at 405 nm (Hoechst), 488 nm (A488, eGFP), 561 nm (tdT, A568), and 639 nm (A647). Multiple beam splitters MBS 488/561/639 and MBS -405 were used to separate fluorescent signals. The software semi-automatically detected brain slices on a slide and adjusted focus. Tile scans of individual brain slices were taken using an AC-Plan-Neofluar 10x/0.3 M27 objective with 0.83 µm x 0.83 µm pixel size, 16-bit depth. Then, individual brain regions containing eGFP-, tdT-, and/or ChAT-positive cells were imaged as Z-stacks. Images were acquired using serial line scan. Z-stacks scanned through the whole slice z range (typically ∼60 µm thickness, 4 µm z-step). The pinhole was adjusted to 1 AU for the longest wavelength fluorophore and fluorescence intensity to below saturation levels. Microscope settings were maintained throughout the experiment for all images for the fluorescent signals with minor changes in gain or laser intensity for the Hoechst 33342 channel.

### Tissue Clearing

After testing several clearing methods, we selected Fast 3D Clear for its consistently reproducible clearing results and compatibility with our computational pipeline for segmenting and quantifying ACh neurons in the mouse brain (Kosmidis et al. (2021)). Brain hemispheres were cleared as previously described (Kosmidis et al. (2021)). All chemicals for tissue clearing were purchased from Merck Millipore Sigma (Burlington, MA, USA). Postfixed mouse brains were cut about 1-2 mm contralateral from the midline to isolate an individual hemisphere. In brief, tissue was repeatedly washed in 0.1 M PBS followed by deionized water. For delipidation, tissue was treated in 50%, 70%, and 90% tetrahydrofuran (THF) in water adjusted to pH 9.0-9.5 with triethylamine at 4°C for 2 hours or overnight, respectively. On the next day, the tissue was rehydrated using 70% and 50% THF in water with triethylamine at 4°C for 2 hours, followed by 100% deionized water overnight, respectively. Finally, brains were immersed in homogeneous clearing solution (48 g histodenz, 10 g urea, 1 g N-methyl D-glucamine, 0.6 g diatrizoic acid, 8 mg sodium azide in 20 ml water, n = 1.515) and incubated on the shaker for at least 24 hours at 37°C for refractive index matching. After reaching full transparency, samples were stored in the fridge.

### Quantitative Image Analysis

Labelling efficiency of neuron populations with tdT and/or eGFP with ChAT was confirmed in most animal models presented in this manuscript. Cells were counted by brain region with the “Point” tool for each channel using ImageJ (RRID:SCR_003070, www.imagej.net). Afterwards, ratios of tdT/ChAT and/or eGFP/ChAT co-positive cells were averaged by brain region.

### Light-Sheet Fluorescence Microscopy

All light-sheet microscopy imaging was performed using a Lightsheet 7 (Zeiss) controlled by the software Zen black (Zeiss). Fluorophores were excited using a 561 nm solid state laser with 9.24 µm light sheet thickness in pivot scan mode for even illumination. The light sheets were generated with two adjustable LSFM 5x/0.1 foc objectives (Zeiss). Fluorescent signals were acquired with an EC Plan-Neofluar 5x/0.16 objective and a PCO Edge 4.2 CMOS camera. Excitation light was filtered using a LBF 405/488/561/640 as main beam splitter and emitted fluorescence was filtered using SBS LP 640 as camera beam splitter and a BP 575-615 emission filter. The objectives were manually adjusted for a centered, thin light sheet in the imaging chamber.

Before imaging, cleared brain hemispheres were transferred into immersion oil (n = 1.515, Cat #: 16482, Cargille labs, Cedar Grove, NJ, USA) and equilibrated for ∼1 hour. Afterwards, the samples were glued onto a sample holder at the caudal cerebellum and transferred into the light-sheet microscope. Then, the sample was immersed in a Meso Chamber (Translucence, Irvine, CA, USA) filled with immersion oil. All samples were imaged in sagittal orientation with the partly missing hemisphere facing the objective. After ensuring the correct sample orientation, the chamber was manually adjusted for the best focus of the light sheet. The focus for both light sheets was then manually aligned using fluorescent cells in a central brain structure (typically caudoputamen, medial/lateral septal nucleus) for orientation. Brains with atypical fluorescence, such as vasculature, other endogenous signals, large air bubbles, or insufficient fluorescence intensity/transparency were discarded.

For volume imaging, the z-range (typically ∼6.5 – 7.5 mm) was defined from the partial contralateral brain hemisphere side to the ipsilateral end of the brain as far as the sample orientation allowed within the chamber, ideally outside the brain or within the most lateral parts of the cortex. Afterwards, the maximal outer boundaries of the brain in sagittal view were used to define the number of tiles (typically 54-72). Tiled z-stacks with 15% overlap were recorded with a voxel size of 0.946 x 0.946 x 2.5 µm (except for CKO66_a3m with 2.0 µm z-step). Both illumination sides were recorded separately to avoid issues with minute light sheet misalignments.

### Image Processing Pipeline

Imaging raw data files were split into individual tiles using Imaris File Converter (Bitplane, Belfast, UK, RRID:SCR_007370). For quality control, one dataset per brain was stitched using Imaris Stitcher and assessed from three angles for structural deformations and distortions, such as an inconsistent midline or blurry cells. Afterwards, tiles of acceptable brains were stitched in Imaris or first converted to tiff in ImageJ using Imaris Reader (https://github.com/PeterBeemiller/ImarisReader) and subsequently stitched using TeraStitcher (https://github.com/abria/TeraStitcher/wiki). The stitched tiff files were downsampled 2×2 (WT, CKO, and CGA) or 3×3 (CVA). After downsampling, images were denoised using custom training data, where multiple noisy and noise-free image pairs were used to train a denoising UNET model (Ronneberger et al. (2015)). A “noise-free” image was generated using average of 500 repetitions of the noisy image (denoising code, https://github.com/SNIR-NIMH/Denoising). Then, the program CATNIP (https://github.com/SNIR-NIMH/CATNIP) was used for cell segmentation and registration to the Allen Brain Reference Atlas v2 (Oh et al. (2014), Bakalar et al. (2023)), and a quantification of the cells and cell volumes by brain regions. To correct for the downsampling, cell pixels were multiplied by 4 and 9 for 2×2 and 3×3 downsampling, respectively. All cell volumes were converted to µm^3^ units.

For the illustrative purposes, a single brain per condition was deconvolved using a recorded point spread function of tdT and subsequently stitched using Huygens Professional 24.10 (RRID:SCR_014237, Scientific Volume Imaging B. V., Hilversum, Netherlands) and rendered in Imaris.

### Quality Control of Cell Segmentation

For the brain regions caudoputamen (CPu), isocortex (ctx), medial habenula (mHb), nucleus accumbens (NAc), and substantia innominata (SI), a visual comparison of fluorescence signals with cell segmentation results at different thresholds was performed to obtain an overall impression of segmentation quality as well as ideal segmentation thresholds. For each animal, the optimal segmentation threshold was then defined based on this evaluation and used to generate region-specific cell counts. To rule out issues with imaging quality, the distribution of cell count per cell volume was compared for left and right illumination side data for the five brain regions. Based on these distribution patterns, minimal cell volume values for these regions were determined as ≥12 pixels (medial habenula) and ≥20 pixels (all other regions), with a maximal cell volume of ≤4000 pixels. These criteria were applied to reduce the number of false-positive puncta, which could occur in the presence of dense innervation or vasculature (see Suppl. Fig. 6 A-E for plots of cell count against cell volume at each threshold). The use of these minimal and maximal cell volume values resulted in similar cell volume distributions for WT, CKO and CGA across segmentation thresholds (median threshold 160k). The maximum cell count of the left and right illumination side of each brain subregion was retained as the cell count for that subregion to account for uneven illumination and decreased fluorescence intensity on the distal side of the brain. Composite regions of interest were defined by pooling anatomically related subregions (e.g., cortical layers, hippocampal subfields) into 108 regions, with each composite region including a minimum of 25 ACh neurons based on WT counts. Major brain regions were defined according to the Allen Reference Atlas Common Coordinate Framework (Wang et al. (2020)).

### Statistical Methods

Analyses were performed in JMP Pro 17.0 (SAS Institute, Cary, NC, USA, RRID:SCR_014242). Statistical significance was set at p ≤ 0.01. To evaluate cell volumes by major brain region, the mean cell volume of each composite brain region was calculated for each animal and then averaged across all composite regions within the corresponding major region. A mixed model was then fit to the log10-transformed major brain region cell volume means as a function of genotype (WT, CKO, and CGA), major brain region, and the interaction between genotype and major brain region, with a random effect to account for within-animal correlation. While a significant difference in estimated cell volumes was detected between different major brain regions, estimated volume differences did not depend on genotype, allowing for detection of ACh neurons of similar volumes across genotypes (Suppl. Fig. 6F). Similar cell counts were seen for major brain regions across segmentation thresholds (Suppl. Fig. 7). Optimal segmentation thresholds for CVA (median threshold 50k) were considerably lower than those for WT, CKO and CGA; thus, CVA cell counts were separately compared to WT cell counts.

To determine similarities between animals, genotypes, and major brain regions, two-way hierarchical clustering of cell counts with Ward linkage was performed, first comparing WT and CKO genotypes, then WT, CGA and CVA genotypes, and finally comparing all four genotypes, with results summarized via heat maps. For each composite region, negative binomial regression models were fit to cell counts as a function of genotype using the log link function and WT as the reference group. A separate negative binomial regression model was fit for CVA due to optimal threshold differences. Regression coefficients were exponentiated to produce rate ratios, a measure of effect size that describes the multiplicative change in cell count relative to WT or other referent; a value of 1 indicated no difference in expected cell counts, values less than 1 indicated lower expected counts, and values greater than 1 indicated higher expected cell counts relative to WT. Due to the large number of comparisons, we controlled for multiple testing using the false discovery rate (FDR) adjustment. FDR-adjusted p-values were transformed into log worth, defined as (-log_10_[FDR(p-value)]), and results were summarized using volcano plots of log worth by estimated rate ratios with composite regions plotted by corresponding major brain region.

## Results

### Loss of GABA co-transmission from ACh neurons does not decrease ACh neuron count in the mouse brain

Co-transmission of ACh and GABA has been described previously in the hippocampus, striatum, mPFC, lateral septum, SNc, as well as the retina (Lee et al. (2010), Saunders et al. (2015a), Saunders et al. (2015b), Estakhr et al. (2017), Sethuramanujam et al. (2016), Takacs et al. (2018), Lozovaya et al. (2018), Obermayer et al. (2019), Granger et al. (2020), Hunt et al. (2022), Le Gratiet et al. (2022), Granger et al. (2023), Lozovaya et al. (2023), Lozovaya et al. (2024)). We recently showed that the loss of GABA co-transmission after conditional knock-out of the vesicular GABA transporter (vGAT, encoded by gene *Slc32a1*) in ACh neurons negatively affects higher brain functions, such as spatial, social, and fear memory as well as habitual learning and locomotion (Goral et al. (2022)). A plausible explanation could be that loss of GABA co-transmission from cholinergic neurons perturbs ACh neuron survival. The lack of ACh signaling could then cause a dementia-like phenotype leading to decreased behavioral performance in spatial, social, and fear memory assays.

To test this hypothesis, we quantified tdT-labelled ACh neurons in vGAT WT and vGAT CKO mice throughout the mouse brain. We previously showed that the ChAT-Cre x Ai14 cross efficiently labels ACh neurons with tdTomato throughout the brain (Goral et al. (2024)). For a more quantitative assessment, we performed tissue clearing followed by light-sheet microscopy to image entire brain at high resolution. When comparing the WT and CKO imaging datasets, we found no notable differences in the gross brain anatomy or the distribution patterns of ACh neurons (Fig. 1 A-B, Suppl. Fig. 3). In single imaging planes, ACh neuron patterns throughout the brain appear comparable between WT and CKO (Fig 1 C-J). In addition, heatmaps of the cell counts for every WT and CKO brain pooled by major brain region (Fig. 2A) and by the 108 composite brain regions (Suppl Fig. 8) indicate no apparent differences between WT and CKO. A volcano plot of estimated rate ratios against log worth values for each of the composite regions did not show significant differences in cell counts between WT and CKO at the p<0.01 level (Fig. 2 B). Taken together, WT and CKO brains are indistinguishable when comparing the brain architecture and ACh neuron distribution patterns or ACh neuron count. We conclude that the loss of GABA co-transmission from ACh neurons does not affect gross brain architecture or ACh neuron distribution patterns in mice.

**Figure 1:**
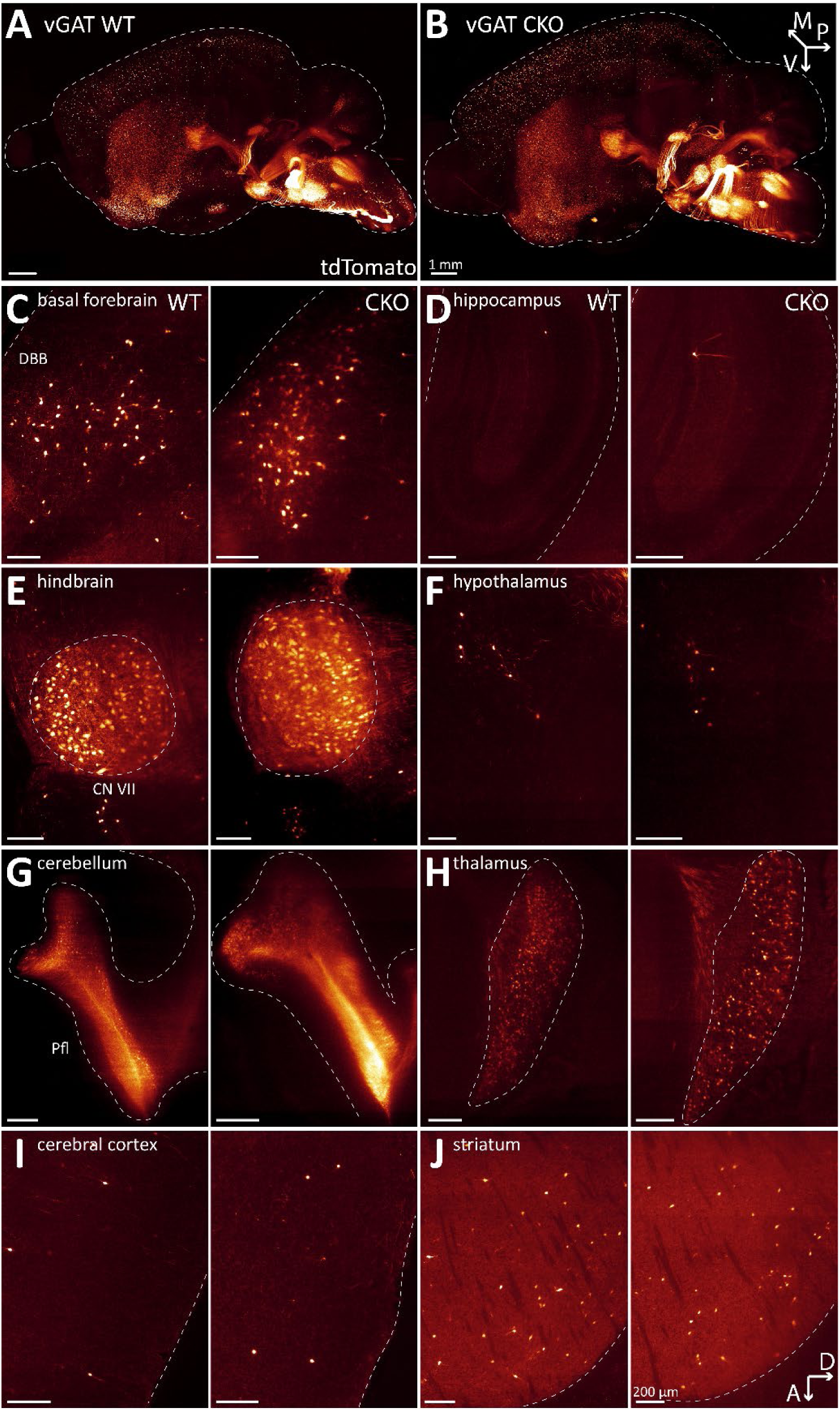
Loss of GABA co-transmission from ACh neurons does not affect distribution of ACh neurons in the whole mouse brain. A-B, Three-dimensional visualization of cleared mouse brain hemispheres at 2 months of age with tdT-labelled ACh neurons for experimental condition vGAT WT (A) and vGAT CKO (B). Arrows indicate orientation of sample: posterior (p, x-axis), ventral (v, y-axis), and medial (m, z-axis). C-J, Depiction of single sagittal imaging planes for vGAT WT (left) and vGAT CKO (right) in basal forebrain (C), hippocampus (D), caudal brain nuclei (E), hypothalamus (F), cerebellum (G), thalamus (H), cerebral cortex (I), and striatum (J). Arrows indicate orientation of imaging planes: dorsal (D, x-axis) and anterior (A, y-axis). Scale bar: 200 µm.

**Figure 2:**
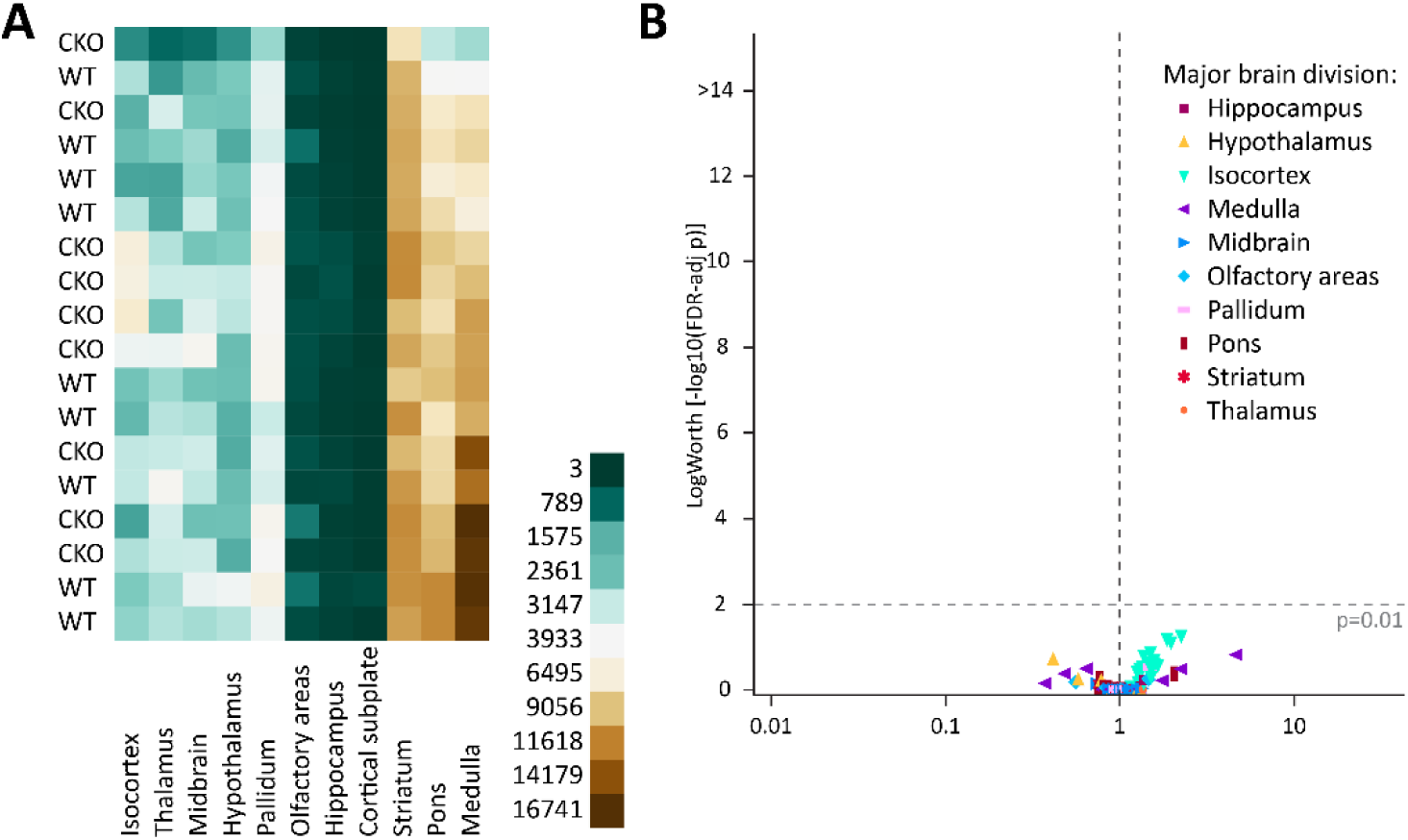
Loss of GABA co-transmission from ACh neurons does not reduce ACh neuron count. A, Heatmap of cell counts of individual WT and CKO brains (rows) by major brain region (columns), with low cell counts in dark green and high cell counts in dark brown. B, Volcano plot of statistical significance (expressed as -log_10_ FDR-adjusted p-value or log worth) by estimated multiplicative change in cell count relative to WT (rate ratio) for 108 composite regions displayed by major brain region. Horizontal dashed line indicates significance threshold (FDR-adjusted p = 0.01); vertical dashed line indicates no change (rate ratio = 1).

### The genetic separation of ACh neurons into Gad2-and vGAT-positive populations

Having ruled out that GABA co-transmission acts by modulating ACh neuron survival, we next set out to systematically assess which ACh neuron populations have a GABAergic fate with various combinations of Cre/FlpO driver and tdT reporter lines (Suppl. Fig. 1, Table 2).

**Table 1:** Antibodies used for Immunohistochemistry.

| Target | Donor<br>Species | Dilution | Vendor | Conjugate | Cat # | RRID |
| --- | --- | --- | --- | --- | --- | --- |
| ChAT | goat | 1:150 | Millipore | N/A | AB144P-200UL | RRID:AB_90661 |
| Goat IgG | donkey | 1:500 | Jackson<br>ImmunoResearch | Alexa<br>Fluor 647 | 705-605-147 | RRID:AB_2340437 |
| RFP | rabbit | 1:500 | Rockland | N/A | 600-401-379 | RRID:AB_2209751 |
| eGFP | chicken | 1:500 | Aves Labs | N/A | GFP-1020 | RRID:AB_10000240 |
| Rabbit<br>IgG | goat | 1:500 | Thermo Fisher | Alexa<br>Fluor 568 | A-11011 | RRID:AB_143157 |
| Chicken<br>IgY | goat | 1:500 | Jackson<br>ImmunoResearch | Alexa<br>Fluor 488 | 103-545-155 | RRID:AB_2337390 |

**Table 2:** Animal model crosses to label ACh/GABA neurons in the brain.

| Targeted neurons | Abbreviation | Cre line | FlpO line | Reporter line |
| --- | --- | --- | --- | --- |
| ChAT positive | WT | ChAT | N/A | Ai14 |
| ChAT positive | CAF | N/A | ChAT | Ai65F |
| ChAT positive | CR | N/A | ChAT | RC::FLTG |
| ChAT/Gad2 positive | CGA | Gad2 | ChAT | Ai65D |
| ChAT/Gad2 positive | CGR | Gad2 | ChAT | RC::FLTG |
| ChAT/vGAT positive | CVA | ChAT | vGAT | Ai65D |
| ChAT/vGAT positive | CVR | ChAT | vGAT | RC::FLTG |

We define GABAergic fate as the initiation of gene expression enabling the neuron to synthesize GABA and express the vesicular GABA transporter (vGAT), thereby possessing the molecular machinery required for GABAergic transmission. In contrast to the cholinergic genes, which are expressed from a single bicistronic gene locus for choline acetyltransferase (ChAT) and vesicular ACh transporter (Slc18a3, encoding the protein vesicular acetylcholine transporter, vAChT), the GABAergic genes for glutamate decarboxylase 1, 2 (Gad1, Gad2), and vGAT are expressed from individual gene loci (Erickson et al. (1994), J J Soghomonian (1998), McIntire (1997)). Consequently, the co-expression of Gad2 and vGAT is consistent with a GABAergic phenotype, that suggests the potential for vesicular GABA release. However, it does not in itself confirm functional GABAergic synaptic transmission.

Due to the availability of suitable animal models, we first had to verify the labeling efficiency of ACh neurons using the ChAT-IRES-FlpO in combination with the FlpO tdT reporter line Ai65F (CAF). Using the CAF model, ACh neurons in forebrain and some midbrain nuclei showed robust ChAT/tdT co-labeling (Suppl. Fig. 2). However, the CAF model was less sensitive targeting ACh neurons in other areas, such as thalamic, some midbrain, and hindbrain nuclei. This confirms that FlpO-based recombination works in principle but is less sensitive in quantitative ACh neuron labeling throughout the brain.

Next, we mapped ChAT/Gad2 neurons using the cross ChAT-FlpO x Gad2-Cre x Ai65D (CGA). In immunostained brain slices, ChAT signals overlapped well with tdT throughout the brain (Fig. 3). In some brain regions, such as the cerebral cortex (ctx), the hippocampus, and the lateral septum (LS), tdT-positive cells were often ChAT-negative or generally had low ChAT expression. In other forebrain nuclei, we detected up to ∼90% ChAT/tdT co-labeling in nuclei of the basal forebrain, such as the medial septum/diagonal band of Broca (MS/NDB). In the caudoputamen (CPu), up to ∼80% ChAT cells were tdT-positive. In the medial habenula (mHb), the abundance of tdT cells was very low or completely absent (not shown). However, mid- and hindbrain nuclei, presented with up to 80% tdT-positive ChAT cells (Fig. 3 C-F). This indicates a strong co-expression of Gad2 in ACh neurons throughout the brain. In addition, it shows that combining Cre and FlpO driver lines increases labeling efficiency compared to a FlpO driver alone.

**Figure 3:**
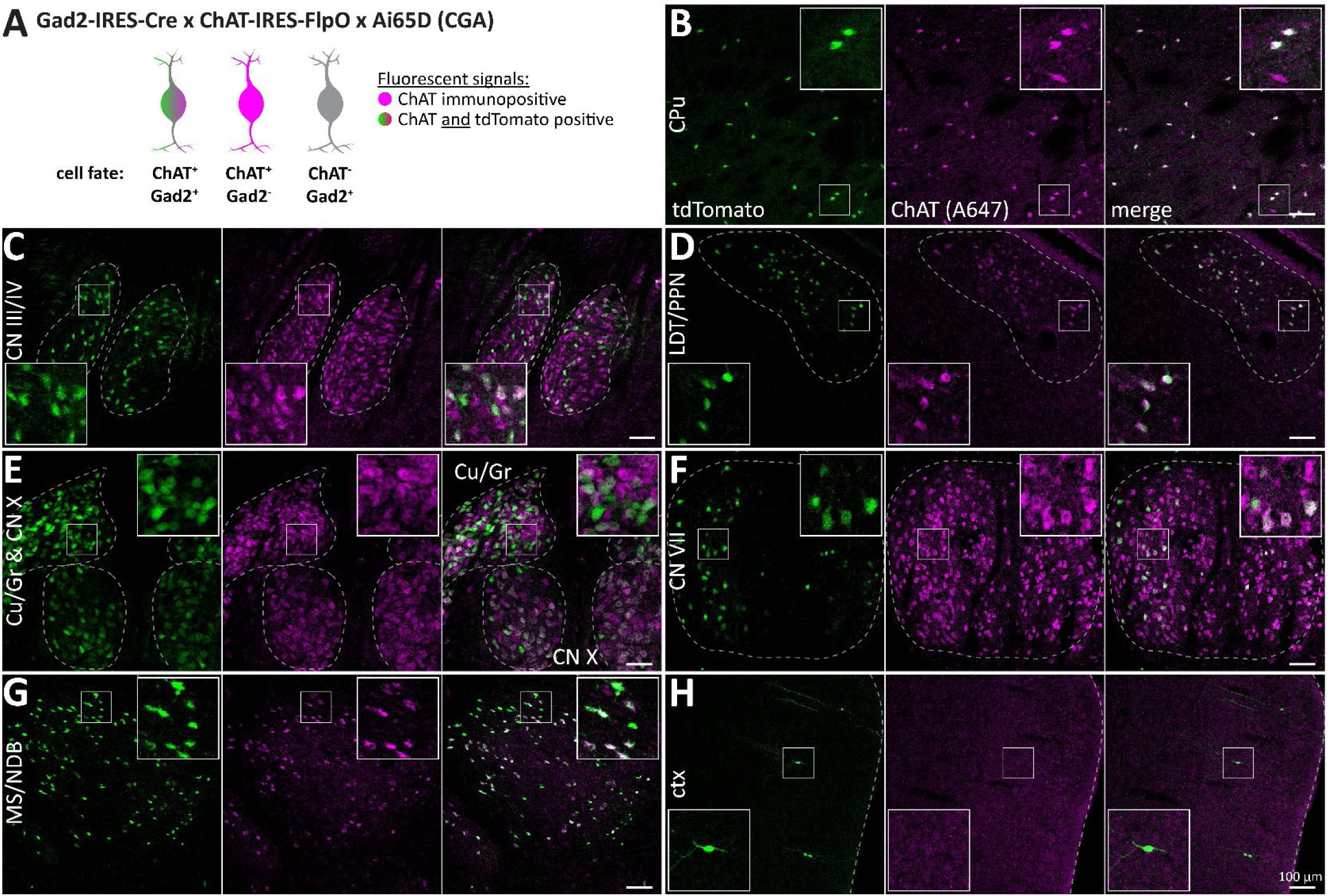
Most forebrain and many mid-/hindbrain ACh neurons are Gad2 positive. A, Animal model used to label cells positive for ChAT and Gad2 with tdT (green and magenta) while neither labeling Gad2-negative ACh neurons (magenta) nor ChAT-negative Gad2 neurons (grey). B-H, 50 µm horizontal brain sections of mice expressing tdT in ChAT/Gad2 neurons were obtained at 2 months of age and counterstained for ChAT. Sections were imaged for tdT (green) and ChAT labelled with A647 (magenta). Maximum intensity projections these brain regions are depicted: caudoputamen (CPu, B), oculomotor/trochlear nerve (CN III/IV, C), laterodorsal tegmentum/pedunculopontine nucleus (LDT/PPN, D), cuneate nucleus/gracile nucleus/vagus nerve (Cu/Gr & CN X, E), facial nerve (CN VII, F), medial septum and diagonal band of Broca (MS/NDB, G), and cerebral cortex (ctx, H). Insets depict magnified ROIs (small white square). Scale bar: 100 µm.

Subsequently, we evaluated the expression of tdT in ChAT/vGAT neurons in slices from animal model ChAT-Cre x vGAT-FlpO x Ai65D (CVA, Fig. 4). We found that ChAT/tdT overlap was decreased compared to the CGA model (Fig. 4). While some brain regions, such as ctx and LS, contained many ChAT-negative tdT-positive cells, high levels of ChAT/tdT co-labeled cells were restricted to MS/NDB as well as oculomotor/trochlear nerves (CN III/IV). In contrast to CGA, in CVA tdT-positive cells were strongly reduced in other regions, such as the nucleus basalis of Meynert/Globus pallidus (MA/GP), the CPu, and throughout the mid- and hindbrain. This indicates that vGAT expression is limited to few ACh neuron nuclei in the mouse brain.

**Figure 4:**
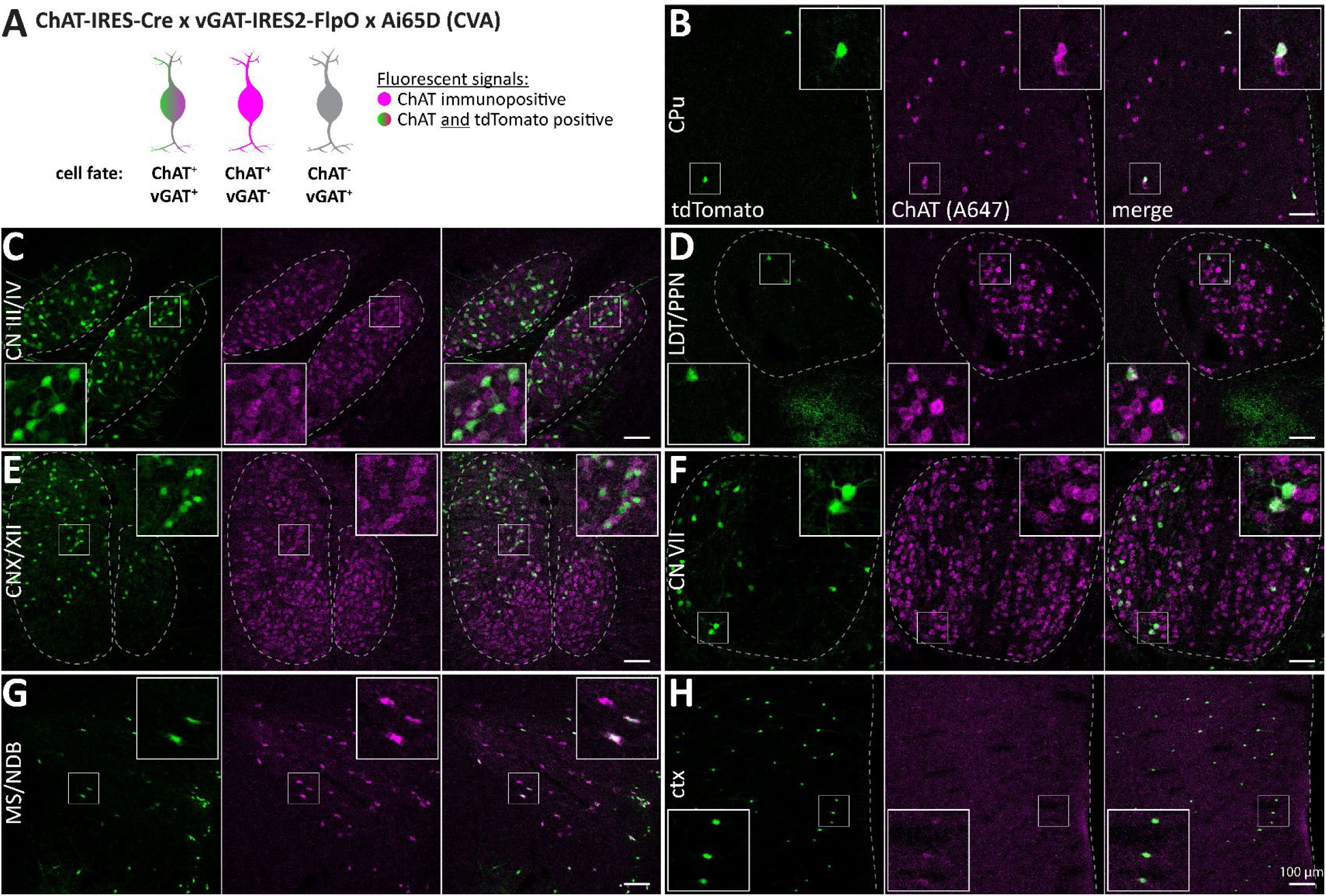
ACh neurons are vGAT positive in few nuclei in the mouse brain. A, Animal model used to label cells positive for ChAT and vGAT with tdT (green and magenta) while not labeling vGAT-negative ACh cells (magenta) nor ChAT-negative vGAT neurons (grey). B-H, 50 µm horizontal brain sections of mice expressing tdT in ChAT/vGAT neurons were obtained at 2 months of age and counterstained for ChAT. Sections were imaged for tdT (green) and ChAT labelled with A647 (magenta). Maximum intensity projections these brain regions are depicted: CPu (B), CN III/IV (C), LDT/PPN (D), vagus/hypoglossal nerve (CN X/XII, E), CN VII (F), MS/NDB (G), and ctx (H). Insets depict magnified ROIs (small white square). Scale bar: 100 µm.

### Many ACh neurons are Gad2-positive, but vGAT expression is far more restricted

To better quantify GABAergic ACh neuron populations, we performed light-sheet microscopy with CGA and CVA brains (Fig. 5, Suppl. Fig. 3-5). The 3D maximum intensity projections indicate that the cholinergic projections leaving brainstem and midbrain toward forebrain or cerebellum are absent or strongly decreased in intensity compared to WT or CKO (Fig 5 A-B). This suggests that the intracranial cholinergic projections may predominantly arise from non-GABAergic ACh neurons. Similarly, fluorescence in cholinergic nuclei within the mid- and hindbrain, such as CN III/IV, V, VI, VII, X, and XII was more diffuse in both CGA and CVA indicating a sparser population of labelled neurons compared to the WT or CKO brains (Suppl. Fig. 3-5). This is consistent with what was previously observed in the histology experiments. In the forebrain, the differences between the CGA and CVA brains require a closer look at individual imaging planes to identify differences. In CVA, the number of tdT neurons in the CPu is strongly reduced (Fig. 5 J). Additionally, when comparing all individual CGA or CVA brains with each other, the forebrain shows a stronger coherence in fluorescence signals, while the mid- and hindbrain signals differ widely between the individual brains (Suppl. Fig. 4-5). Therefore, the pattern of GABAergic marker co-expression or transgene expression within midbrain and hindbrain ACh neurons appears relatively diffuse and variable, whereas in the forebrain it is more consistently regulated. This may stem from generally low or heterogeneous levels of GABAergic marker expression in mid- and hindbrain regions which may be insufficient to reliably drive recombination across those areas.

**Figure 5:**
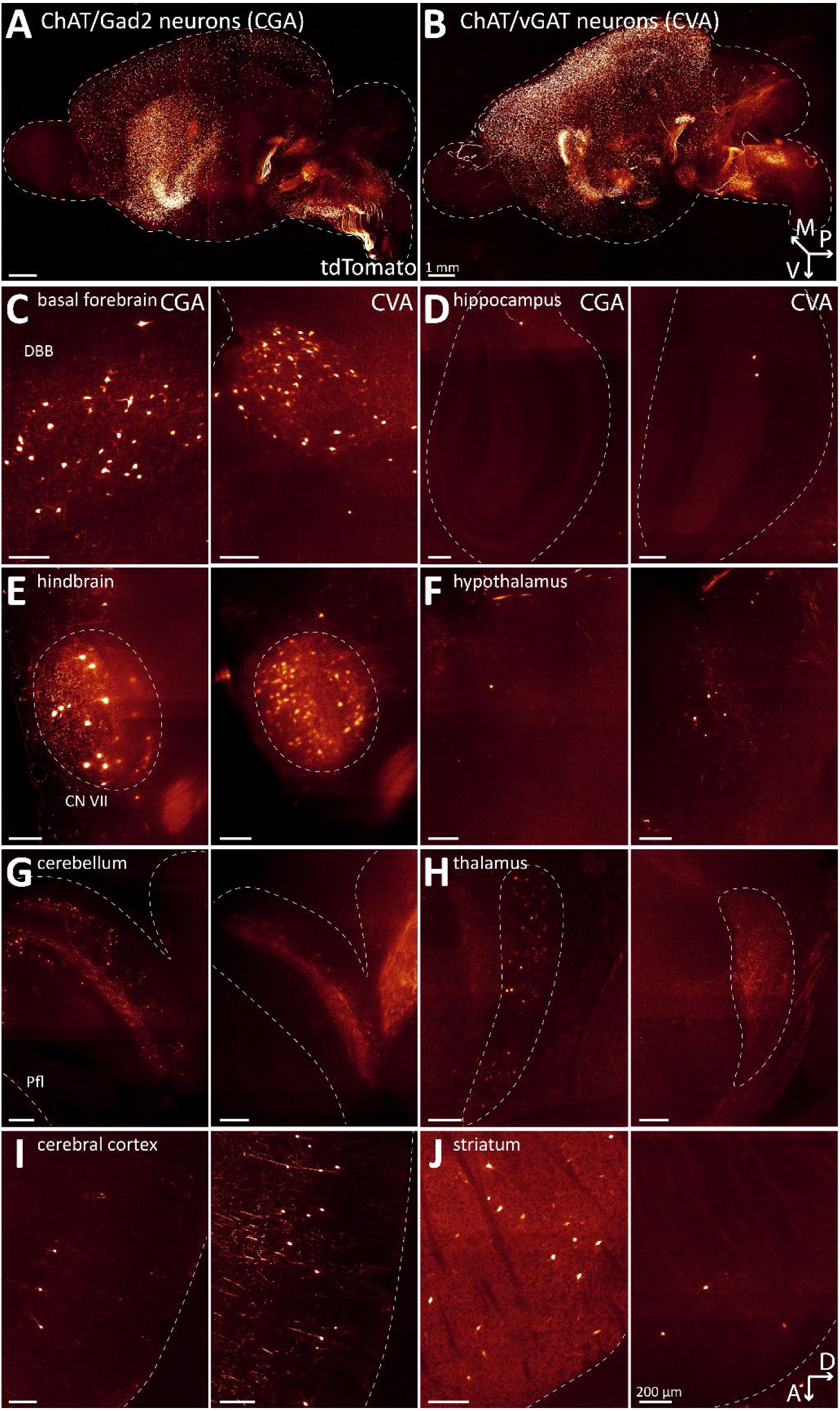
Distribution of patterns of ChAT/Gad2 neurons and ChAT/vGAT neuons in the whole mouse brain. A-B, Three-dimensional visualization of ChAT/Gad2 neurons (A) and ChAT/vGAT neurons (B) in cleared mouse brain hemispheres at 2 months of age. Arrows indicate orientation of sample: posterior (p, x-axis), ventral (v, y-axis), and medial (m, z-axis). C-J, Depiction of single sagittal imaging planes for CGA (left) and CVA (right) in basal forebrain (C), hippocampus (D), caudal brain nuclei (E), hypothalamus (F), cerebellum (G), thalamus (H), cerebral cortex (I), and striatum (J). Arrows indicate orientation of imaging planes: dorsal (D, x-axis) and anterior (A, y-axis). Scale bar: 200 µm.

Hierarchical clustering of cell counts by major brain region showed that WT and CGA brains clustered together, while CVA brains formed a separate cluster (Fig. 6 A). This pattern was also observed for hierarchical clustering of cell count by composite brain region for the four genotypes (Suppl. Fig. 8). Cell count ratios of CGA relative to WT indicated that Gad2-positive cell count in the CGA group is mostly reduced in composite regions of the thalamus, and subregions of hypothalamus (e. g. posterior hypothalamic nucleus (PH), supramammillary nucleus (SUM)), midbrain (e. g. ventral tegmental area (VTA), interpeduncular nucleus (IPN)), medulla (nucleus raphe magnus (RM), paragigantocellular reticular nucleus (PGRN), magnocellular reticular nucleus (MARN)), and pons (tegmental reticular nucleus (TRN), nucleus of the lateral lemnicscus (NLL), pontine reticular nucleus (PRNr)) (Fig. 6 B). While there is also a significant cell count reduction for CGA relative to WT in nuclei of striatum and pallidum, the decrease is much smaller (ranged between 10-30%). Isocortex, LS, and hippocampus do not present with a reduction in cell count in CGA indicating that ACh neuron labeling in the forebrain is comparable despite the use of both different driver and reporter lines.

**Figure 6:**
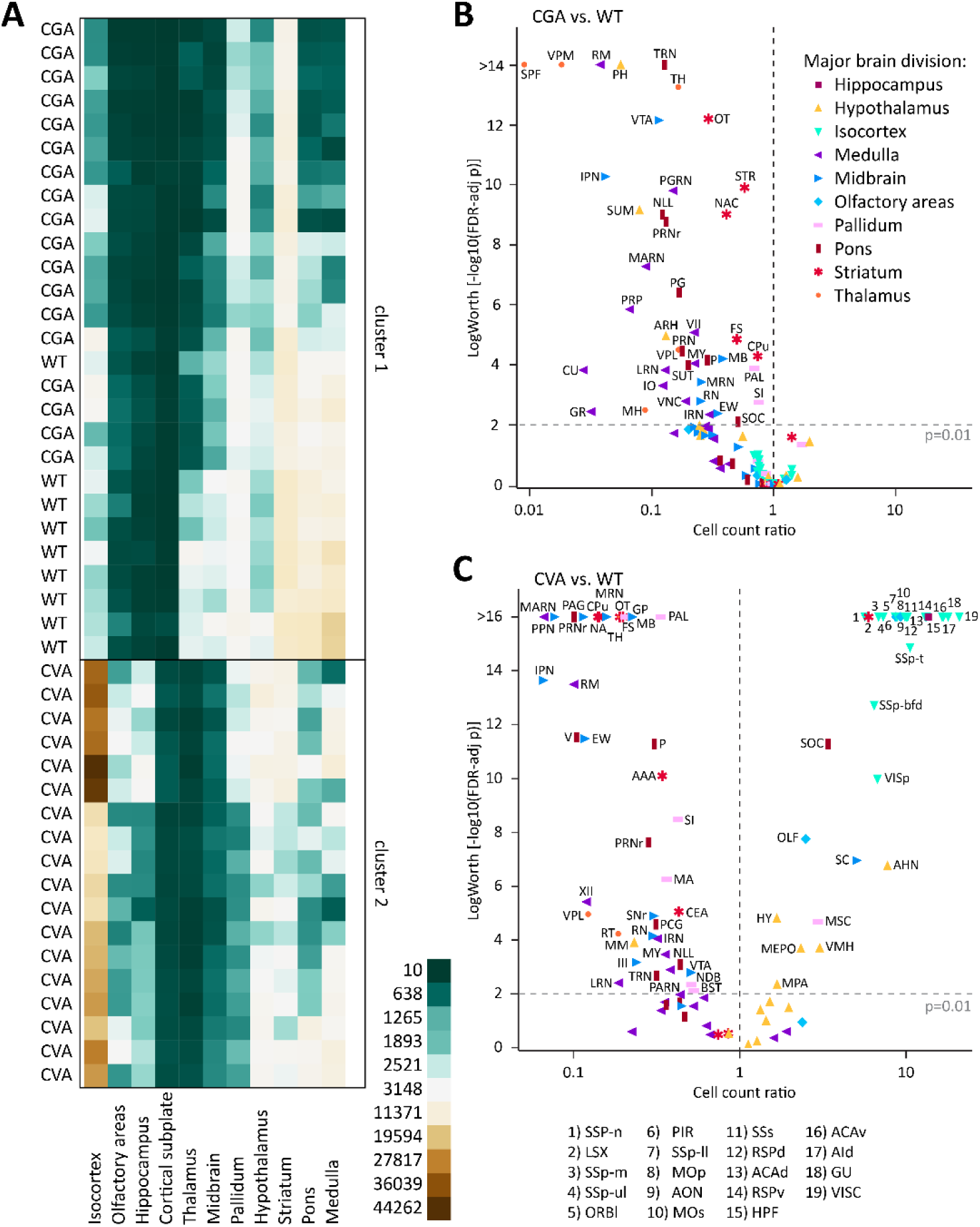
ChAT/Gad2 and ChAT/vGAT overlap is limited to few nuclei in the brain. A, Heatmap of cell counts of individual animal WT, CGA, and CVA brains (rows) by major brain region (columns), with low cell counts in dark green and high cell counts in dark brown. B-C, Volcano plot of statistical significance (expressed as -log_10_ FDR-adjusted p-value or log worth) by estimated multiplicative change in cell count for CGA (B) and CVA (C) relative to WT (rate ratio) for 108 composite regions displayed by major brain region. Horizontal dashed line indicates significance threshold (FDR-adjusted p = 0.01); vertical dashed line indicates no change (rate ratio = 1).

In contrast, cell count reductions in CVA relative to WT were predominantly detected in striatum, midbrain, pons, and medulla (Fig. 6 C). However, there were also reductions in some composite regions of the pallidum, thalamus, and hypothalamus. Reductions in typical ACh neuron composite regions, such as the CPu, NAc, SI, and MA were more dramatic in CVA than in CGA (CP: 86% vs 26%, NA: 86% vs 60%, SI: 58% vs 25%, MA: 64% vs 7%), while reductions in NDB were moderate (49% vs 26%). The MSC showed 2.95x increase in cell counts (p<0.0001) relative to WT. In contrast to CGA, we detected many composite regions in the CVA brains with significantly increased cell counts, ranging from 5-20x higher than WT. These increases were predominantly seen in nuclei of the isocortex, lateral septal complex (LSX), hypothalamus, and hippocampal formation (HPF). This indicates potential off-target effects associated with one or several of the animal lines. We did not find sex differences in any of the composite brain regions for CGA or CVA brains (Suppl. Fig. 9 B, D).

Taken together, this indicates that labeling of ACh/GABA co-positive neurons strongly depends on the individual brain composite region. In addition, the combination of different driver lines has a strong impact on labeling throughout the brain.

### The distribution of Gad2- and vGAT-positive ACh neuron populations is robust independent of reporter line

To further assess potential off-target effects of the transgenic mouse lines, we verified labeling efficiency using the Cre/FlpO dual reporter line, RC::FLTG. The benefit of this line is the separate expression of tdT in FlpO positive cells and eGFP in FlpO/Cre co-positive cells (Plummer et al. (2015)). We first used this animal model to characterize ACh neuron labeling efficiency using cross ChAT-FlpO x RC::FLTG (CR, Suppl. Fig. 10). We found that the overall labeling efficiency of ACh neurons was reduced compared to CAF indicating that the RC::FLTG reporter is less sensitive to FlpO expression than reporter line Ai65F. We assumed that this might reduce potential off-target effects of the driver lines and may yield some insights into the expression levels of GABAergic markers in ACh neurons.

To confirm the distribution of ChAT/Gad2 co-positive neurons, we used the cross ChAT-FlpO x Gad2-Cre x RC::FLTG (CGR, Fig. 7 A left). Generally, the labeling efficiency of ChAT/Gad2 neurons in model CGR was decreased compared to both CAF and CGA. However, we found that CGR labels more than 50% of ChAT neurons in the forebrain with GFP. This includes cells in basal forebrain, striatum, cerebral cortex. In the midbrain, we detected GFP-positive cells predominantly in CN III/IV. In the hindbrain the distribution of GFP-positive cells was negligibly small (<10% of ChAT cells). This indicates a generally low Gad2 expression throughout the hindbrain. Likewise, tdT-positive ChAT cells were low compared to CAF, with the highest tdT/ChAT ratio in the laterodorsal tegmentum/pedunculopontine nucleus (LDT/PPN). This left up to 50% of ChAT cells unlabeled throughout the brain. Taken together, we observed some degree of insensitivity of the RC::FLTG reporter line if the cells are only expressing FlpO. However, FlpO/Cre co-expressing cells showed improved labeling efficiency compared to the FlpO-only cross, CR.

**Figure 7:**
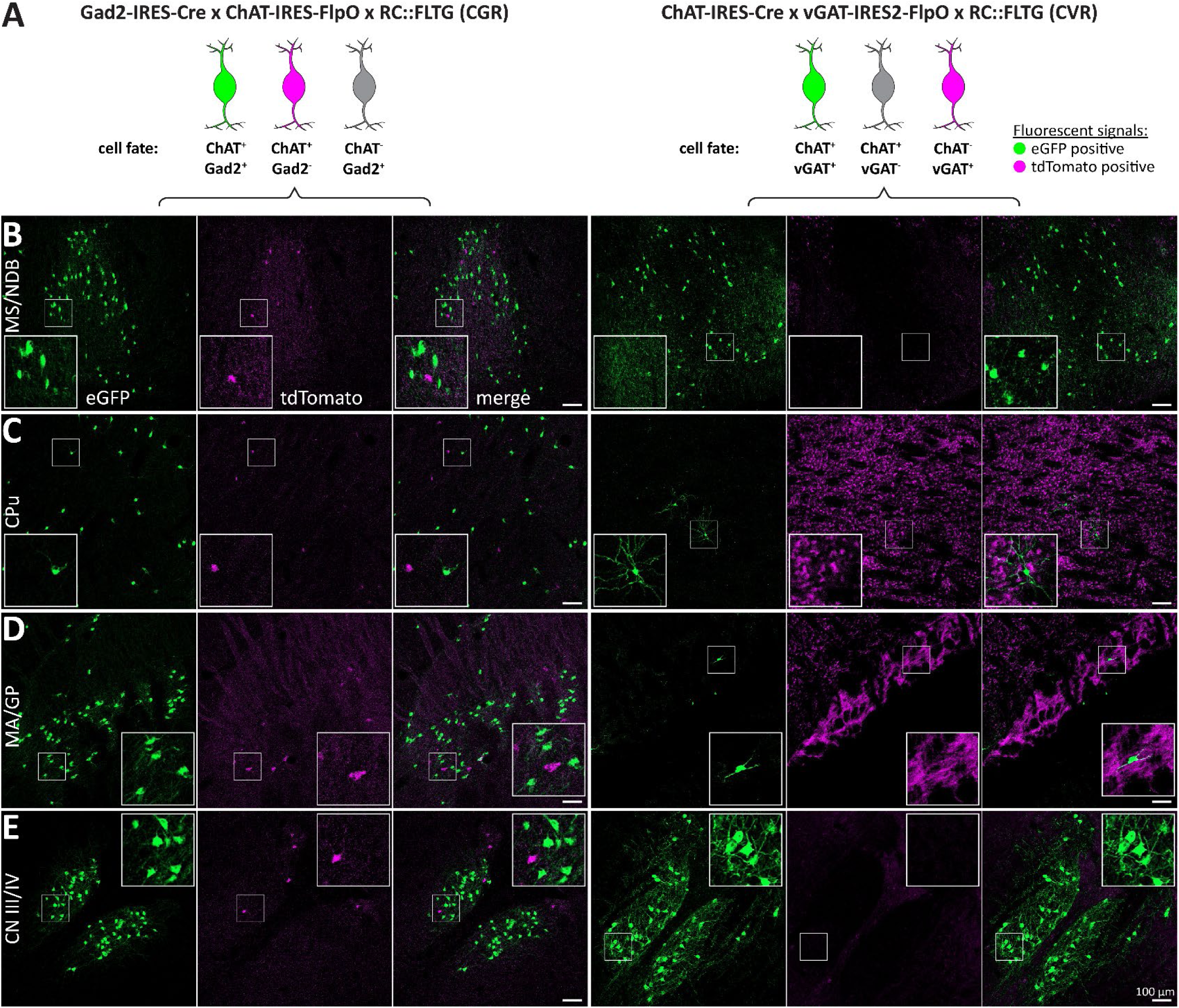
The distribution of Gad2- and vGAT-positive ACh neuron populations is reproducible using a different reporter line. A, Left: Animal model used to label ChAT/Gad2 cells with eGFP (green) and ChAT cells with tdT (magenta), while not labeling ChAT-negative cells (grey). Right: Animal model to label ChAT/vGAT cells with eGFP (green) and ChAT cells with tdT (magenta), while not labeling vGAT-negative cells. Slices were stained for tdT and eGFP to enhance fluorescent signals. B-H, 50 µm horizontal brain sections of mice were obtained at 2 months of age. Sections were imaged for eGFP (green) and tdT (magenta). These brain regions are depicted: MS/NDB (B), CPu (C), MA/GP (D), and CN III/IV (E). Insets depict magnified ROIs (small white square). Scale bar: 100 µm.

Next, we confirmed the distribution of ChAT/vGAT co-positive neurons by using the cross ChAT-Cre x vGAT-FlpO x RC::FLTG (CVR, Fig. 7 A right). We found that model CVR labels similarly to CVA, with highly abundant GFP cells in the cortex, MS/NDB, and some cells in CN III/IV. Comparable to CVA, GFP-positive cell count is very low in both MA/GP and CPu confirming previous findings. In the hindbrain, GFP-positive cell numbers are very low, with the highest number in CN X/XII. We detected no tdT-positive ChAT cells in CVR indicating that Cre expression and recombination is not the limiting factor in labeling efficiency. However, tdT expression in non-ChAT cells and fibers is high throughout the brain. Specifically, we found widespread tdT labeling of cells and fibers in the cortex, the striatum, the globus pallidus, the indirect pathway projections to the midbrain, fibers within the cerebellar arbor vitae, and the Purkinje layer. This indicates that the vGAT-FlpO line in combination with the RC::FLTG reporter is sufficient to label GABAergic neurons in the brain. The use of the RC::FLTG reporter line corroborates our previous findings, demonstrating that ChAT/Gad2 neurons are more numerous across the mouse brain than ChAT/vGAT positive neurons.

### Gad2/vGAT co-positive ACh neurons are restricted to few nuclei in the brain

To evaluate the coherence of the experiments to co-label GABAergic ACh neurons with ChAT, we pooled the fluorophore/ChAT ratios by brain region and genotype (Fig. 8). All animal crosses which are supposed to label all ACh neurons show various degrees of labeling efficiency with the highest efficiency in WT, while both CAF and CR are less efficient than WT (Fig. 8 A-C). In addition, most nuclei in non-WT crosses show higher variability compared to WT. Interestingly, the dual recombinase crosses CGA and CVA show increased labeling efficiency compared to their respective FlpO-only controls (Fig. 8 D-G). To interpret the overlap of the different subpopulation of ACh neurons, we classified mean labeling efficiency in four different bins (>10%, 10-50%, 50-75%, and 75-100%). When comparing the conditions WT, CGA, CGR, CVA, and CVR based on this classification, we found a high consistence in distribution between ChAT/Gad2 and ChAT/vGAT neurons only in the nuclei MS/NDB with at least 50% co-labeling, and CN III/IV with at least 25% co-labeling. Other regions, such as MA/GP or CPu, present with a co-labeling efficiency <10% for at least one cross. In addition, there are low ChAT-expressing cell populations in the cerebral cortex, hippocampus, and LS that were consistently labelled indicating a dual ACh/GABA character. Our data suggests that ACh neuron populations with the potential to quantitatively co-transmit GABA are restricted to few nuclei in the mouse brain.

**Figure 8:**
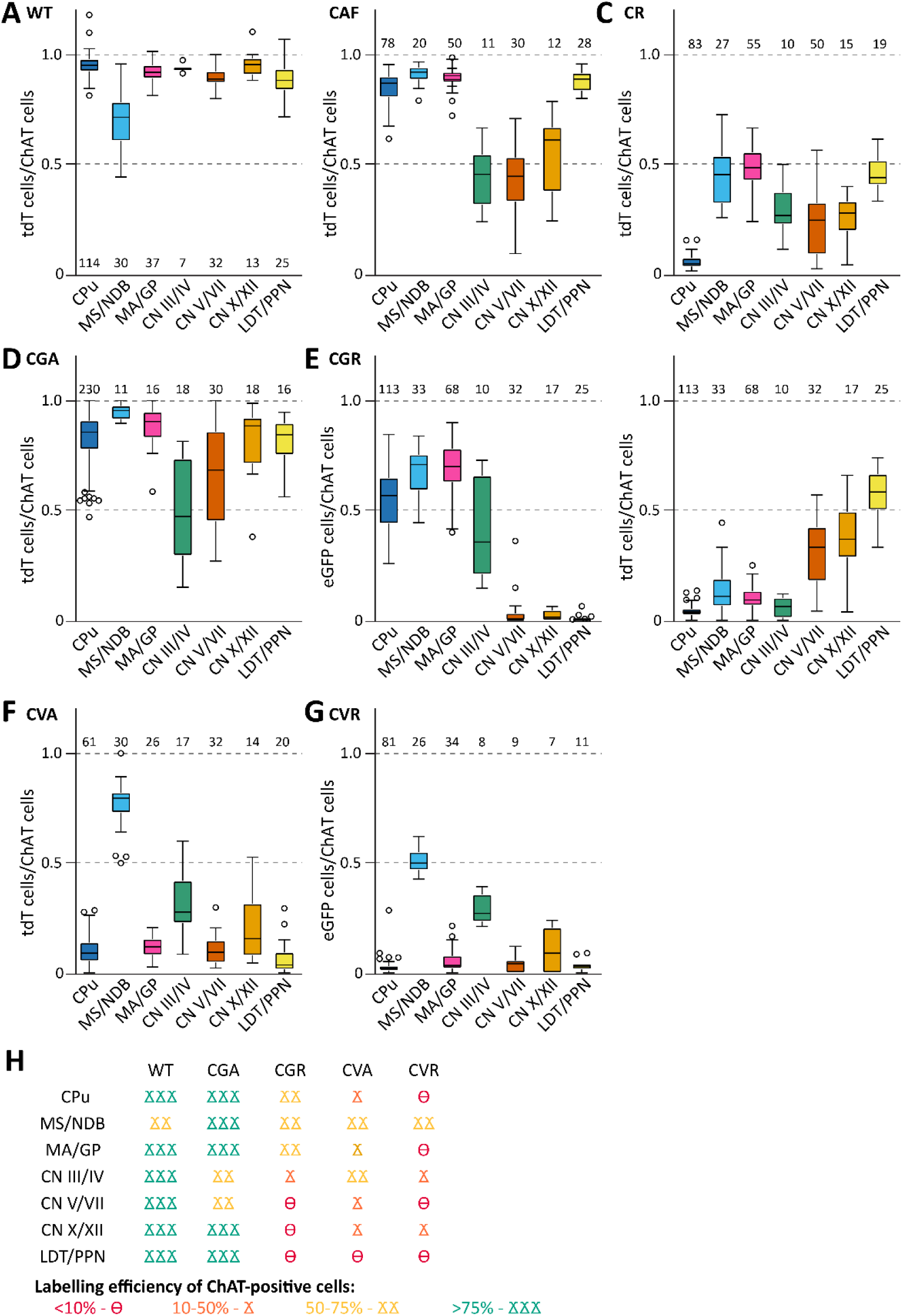
Labeling efficiency of cholinergic neurons is dependent on combination of driver and reporter lines. A, Labeling efficiency of ChAT-positive neurons with tdT in WT brain slices. Part of this data was previously published Goral et al. (2024). B, Labeling efficiency of ChAT-positive neurons with tdT in CAF brain slices. C, Labeling efficiency of ChAT-positive neurons with tdT in CR brain slices. D, Labeling efficiency of ChAT-positive neurons with tdT in CGA brain slices. E, Labeling efficiency of ChAT-positive neurons with eGFP (left) and tdT (right) in CGR brain slices. F, Labeling efficiency of ChAT-positive neurons with tdT in CVA brain slices. G, Labeling efficiency of ChAT-positive neurons with eGFP (left) and tdT (right) in CVR brain slices. H, summary cartoon of labeling efficiency of ChAT-positive neurons. Boxes indicate interquartile range (25^th^-75^th^ percentiles); whiskers extend to minimum and maximum values (excluding outliers) and outliers as gray circles. Numbers of individual data points contributing to the distribution are written over or underneath each boxplot. At least 3 biological replicates including both males and females were used for every experiment.

## Discussion

We present mesoscopic scale datasets of various ACh neuron populations across the mouse brain, yielding these key insights: 1) ACh neuron counts remain unchanged in the absence of GABA co-transmission; 2) ACh/Gad2 neurons are numerous in both the fore- and hindbrain; and 3) ACh/vGAT neurons are confined to a limited subset of nuclei. By combining histology and light-sheet microscopy of cleared brain hemispheres, we studied the distribution and probed the fate of ACh neurons in transgenic mice. Our data provides new insights into the heterogeneity of ACh neurons and shed light on the significance of GABA co-transmission in ACh neuron function.

### ACh/GABA neuron populations in the brain

Co-transmission of GABA from ACh neurons has been reported in several brain regions and is associated with behavioral performance in rodents (Saunders et al. (2015a), Saunders et al. (2015b), Estakhr et al. (2017), Takacs et al. (2018), Lozovaya et al. (2018), Obermayer et al. (2019), Granger et al. (2020), Le Gratiet et al. (2022), Goral et al. (2022)). Our findings indicate that the loss of GABA co-transmission does not reduce ACh neuron count throughout the brain. This suggests that perturbed GABA co-transmission affects higher brain functions either during neural circuit formation or acute neuromodulation effects, but not through a loss of cholinergic signaling. However, more work is needed to systematically assess when GABA co-transmission plays its most critical functions. Recently, the developmental role of GABAergic marker expression in ACh neurons has become an emerging topic (Granger et al. (2023), Lozovaya et al. (2023), Lozovaya et al. (2024)). In the hindbrain of chick embryos, it has been reported that some ACh neurons express GABAergic markers potentially supporting a role of GABA during development (von Bartheld (1989)). In addition, there are reports on neurotransmitter switch from GABA to ACh or vice versa during development in parts of the basal forebrain, including the MA, the MS/NDB, and the LS (Hunt et al. (2022), Granger et al. (2023)). This is conducive of a separate role of GABA co-transmission and regulation of cholinergic and GABAergic genes depending on the developmental stage of the organism and brain region.

During brain formation, ACh neurons originate from lineages and proliferation zones that also give rise to GABAergic interneurons (Ahmed et al. (2019), Allaway et al. (2020), Ananth et al. (2023)). In the mouse CPu, about half of the ACh neurons are Lhx6 positive and most contain Gad2 mRNA (Lozovaya et al. (2018)). This concurs with our observation that most striatal ACh neurons were tdT positive in our ChAT-FlpO x Gad2-Cre crosses. However, we found most striatal ACh neurons are vGAT negative judging by the reporter line fluorophore expression patterns. This indicates that striatal ACh neurons are either predominantly precluded from co-transmitting GABA or the vGAT expression is insufficient to drive the FlpO-mediated recombination and activate expression of the transgenic reporter fluorophore.

In the basal forebrain, there is a dichotomy in distribution between MS/NDB with a high abundance of Gad2/vGAT co-positive ACh neurons, but a low abundance of Gad2/vGAT co-positive ACh neurons in the more caudal regions, such as MA and SI. MS/NDB ACh neurons originate predominantly from later born Nkx2.1/Zic4 lineages from the septal epithelium, while MA, SI, and VP are mostly non-Zic4 lineages from the medial ganglionic eminence and the ventral pallium (Ananth et al. (2023)). In addition to cell lineage origin, the organization of BF projections can be approximately separated between MS/NDB and the more caudal BF nuclei (Varsanyi et al. (2025)). It is possible that distinct lineages may differentially regulate vGAT expression in these neurons (Kilpinen et al. (2023)). Our data presented here supports the assumption that the GABAergic genes Gad2 and Slc32a1 are regulated independent of each other in certain cell types. Further research is required to disseminate how GABAergic genes are regulated throughout the brain and what non-canonic roles GABAergic genes fulfill in addition to neurotransmission in different cell populations.

### What makes a neuron GABAergic?

The regulation of GABAergic neuron subtype development is highly complex (Lim et al. (2018)). The promoter regions for the GABAergic genes Gad1/2 and Slc32a1 show little cell-type specificity when used in viral constructs (Duba-Kiss et al. (2021)). Tissue-specific expression of GABAergic markers likely depends on additional regulatory elements, such as enhancers (Kilpinen et al. (2023)). Gad1, Gad2, and vGAT, are often treated interchangeably, but they are distinct genes with different functions and expression patterns (Enoch et al. (2013), Asada H (1996), M Esclapez (1994), Walls et al. (2010)). Considering that especially Gad1/2 have been implied in fulfilling other functions besides providing GABA for neurotransmitter release, it may not be surprising that GABAergic genes are regulated independently (Le et al. (2017)). GABAergic synapse markers, such as gephyrin and neuroligin 2, can also be found in dopaminergic or cholinergic synapses (Uchigashima et al. (2016), Takacs et al. (2013), Takacs et al. (2018)). In addition, midbrain dopaminergic neurons release GABA after membrane uptake, but these neurons do not synthesize GABA intracellularly (Tritsch et al. (2014), Berrios et al. (2016)). Likewise, it has been shown that the interactome of nicotinic ACh receptors includes GABAergic receptors (Rosenthal et al. (2025)). Given that GABA co-transmission has been reported with faster kinetics than ACh release, a specialized active zone organization is required for the independent release of GABA (Lee et al. (2010), Takács (2018)). This shows that GABAergic gene regulation in non-canonical GABAergic neurons is flexible, and postsynaptic structures provide the framework for integrating GABA co-transmission into synapses.

Further study is needed on how the heterogeneity of cholinergic and GABAergic fate is regulated during development (Su-Feher et al. (2022), Li et al. (2025), Bright et al. (2025)). Specifically, subsets of cortical GABAergic VIP neurons express cholinergic markers, while cholinergic gene expression is absent in other GABAergic neuron lineages or changes during brain maturation (Li et al. (2018), Dudai et al. (2020), Granger et al. (2020), Hunt et al. (2022)). It is also important to examine the molecular profiles of ACh and ACh/GABA neurons in brain and spinal cord need to be assessed at different time points to better quantify GABAergic gene expression and GABA co-transmission in ACh neurons throughout development.

### Experimental limitations of this study

Our study relied on transgenic animal lines that combine Cre and FlpO recombinase activity to drive the expression of fluorescent marker expression in subsets of ACh neuron populations. This approach has been successfully employed before to efficiently label cholinergic neuron populations (von Engelhardt et al. (2007), Saunders et al. (2015a), Li et al. (2018), Allaway et al. (2020), Goral et al. (2024)). However, unspecific labeling and other off-target effects associated with recombinase activity have been previously reported (Nasirova et al. (2020)). Our own data presented here indicates that not every recombinase is equally efficient to drive fluorescent transgene expression. Specifically, we found insufficient labeling of cholinergic neurons using the ChAT-IRES-FlpO in combination with two reporter lines, Ai65F and RC::FLTG, in some brain regions contrasting previous experiments using a ChAT-Cre driver line (Goral et al. (2024), Li et al. (2018)). Surprisingly, for the ChAT-FlpO x Gad2-Cre crosses, fluorescent marker expression was substantially improved, particularly in the forebrain. This suggests that low Flp expression is likely a limiting factor for efficient labeling consistent with previous reports (Werdien D. (2001), Zhao et al. (2023)). In contrast, high Flp expression is more efficient for recombination of transgenes than Cre suggesting cell-type specific differences in labeling efficiency (Takata et al. (2011)). Since vGAT expression in basal forebrain ACh neurons was reported low compared to Gad1/2 expression, this might contribute to discrepancy in our vGAT-FlpO x ChAT-Cre dataset from previous reports (Granger et al. (2023)).

For practical reasons, we glued the sample holder to the caudal cerebellum of the cleared brain. This obstructed the light path of some of the caudal brainstem, preventing the detection of cells within some cholinergic regions, such as CN X, CN XII, nucleus ambiguus, and the spinal cord. Our histology experiments, however, indicate that CN X/XII contain Gad2/ChAT and ChAT/vAChT cell counts at levels comparable to more anterior nuclei in the hindbrain.

### Future directions

More research is required to better understand co-release and co-transmission in general. In particular, additional studies are required on the role of protons, nucleic acids, and other neurotransmitters (Upmanyu et al. (2022), Burnstock (1976), Nelson et al. (2014)). One of the main reasons that precludes an in-depth study of heterogeneity in ACh neuron subpopulations is the sparsity within many brain regions, which yield few data points in single-cell sequencing experiments (Munoz-Manchado et al. (2018), Garma et al. (2024)). Likewise, techniques, such as bulk RNAseq or Spatial Transcriptomics dilute the individual differences between ACh neurons. Future research also needs to address ACh neuron heterogeneity regarding input/output relationships of individual ACh neurons, cell morphology, circuit function, and projection patterns (Gielow and Zaborszky (2017), Alkaslasi et al. (2021), Yayon et al. (2023), Kim et al. (2024), Varsanyi et al. (2025)).

## Supporting information

SuppFile_Goral_et_al

## Acknowledgments

This research was supported by the Intramural Research Program of the National Institutes of Health (NIH). The contributions of the NIH authors are considered Works of the United States Government. The findings and conclusions presented in this paper are those of the authors and do not necessarily reflect the views of the NIH or the U.S. Department of Health and Human Services. We are grateful for the NIEHS Comparative Medicine Branch (CMB) Animal Resources Section for providing general animal housing and care as well as the NIEHS CMB Veterinary Medicine Section for providing general veterinary care. Specifically, animal care was provided by Scotty D. Dowdy, Maria G. Barrientos Sandoval, Alexis D. Wagner, Robert Vaquiz, and Iz’Jay M. Sessoms. We thank Erica L. Scappini, Christopher R. Day, Dr. Lenka Radonova, and C. Jeff Tucker of the NIEHS Fluorescence Microscopy & Imaging Center (FMIC) for technical assistance. All samples that contribute to the datasets in this manuscript were imaged with hardware provided by the NIEHS FMIC. Initial light-sheet data handling was performed using FMIC computer hardware. We thank Dr. Pablo Ariel at the Microscopy Services Laboratory, University of North Carolina, Chapel Hill for usage of the light-sheet microscope for pilot experiments. We thank Dr. Harshad Vishwasrao of the Advanced Imaging Resource at NIBIB for the usage of the DISPIM for pilot experiments. IT assistance for large data handling was provided by Christopher L. Stone, John D. Grovenstein, Kenneth Grantham, Frank S. Day, Adam B. Burkholder, as well as Drs. David C. Fargo and Charles P. Schmitt. This work utilized the computational resources of the NIH High Performance Computing Biowulf cluster (HPC, http://hpc.nih.gov). We thank Caroll A. Co and Dr. Sandra J. McBride for statistical support funded by the NIEHS under contract to Social and Scientific Systems, Inc., A DLH Holdings Corp Company (contract #: GS-00F-173CA / 75N96022F00055). We thank James L. Hunnicutt Jr. and the Fabrication and Repair Studio for designing and making specialized pieces for light-sheet microscopy. Color palettes suitable for color blind individuals were chosen according to recommendations by Dr. Martin Krzywinski (https://mk.bcgsc.ca/color/). We thank Drs. Shiyi Wang, Robert P. Machold, Lorna W. Role, Pantelis Tsoulfas, Patricia Jensen, Nick W. Plummer, and members of the Yakel lab for valuable discussions or comments on experiments or the manuscript.

