## Supplementary material for "Mesoscopic analysis of GABAergic marker expression in acetylcholine neurons in the whole mouse brain": SuppFile_Goral_et_al

**Supplemental Figures**

**Suppl. Figure 1: Breeding strategy of mouse lines to label specific ACh neuron populations in the brain.**

A, ACh neurons were labelled with tdT by crossing ChAT-IRES-Cre mice with Ai14 Cre reporter mice. The offspring was used to conduct light-sheet microscopy with cleared brains. Refers to data presented in Fig. 1-2 and Suppl. Fig. 3, 8. This cross is considered the “WT” control condition and was previously characterized to efficiently label ACh neurons throughout the brain (Goral et al. (2024)). B, Brains lacking GABA co-transmission from ACh neurons were generated using a conditional knock-out for vGAT. All ACh neurons were labelled with tdT. The offspring was used to conduct light-sheet microscopy imaging. Data presented in Fig. 1-2 and Suppl. Fig. 3, 8. This cross is considered “vGAT CKO”. C, ACh neuron populations were genetically divided into a GABAergic subpopulation, Gad2-IRES-Cre mice were crossed with ChAT-IRES-FlpO and Ai65D Cre, FlpO reporter mice. Only cells that expressed at one point ChAT and Gad65 were tdT positive. The offspring was used to conduct light-sheet microscopy imaging and immunohistochemistry with brain slices. Data presented in Fig. 3, 5, 6, and Suppl Fig. 4, 8-9. This cross is considered “CGA”. D, ACh neuron populations were genetically divided into a GABAergic subpopulation, ChAT-IRES-Cre mice were crossed with vGAT-IRES2-FlpO-D and Ai65D Cre, FlpO reporter mice. Only cells that expressed at one point ChAT and vGAT were tdT positive. The offspring was used to conduct light-sheet microscopy imaging and immunohistochemistry with brain slices. Data presented in Fig. 4-6 and Suppl Fig. 5, 8-9. This cross is considered “CVA”. E, To verify labeling efficiency of the ChAT-IRES-FlpO driver line, ACh neuron populations were genetically labelled using ChAT-IRES-FlpO crossed with Ai65F FlpO reporter mice. The offspring was used to conduct light-sheet microscopy imaging and immunohistochemistry with brain slices. Data presented in Suppl. Fig. 2. This cross is considered “CAF”. F, To further verify labeling efficiency of ChAT-IRES-FlpO with a different reporter line, ChAT-IRES-FlpO was crossed with RC::FLTG FlpO reporter mice. Cells that expressed at one point ChAT were tdT positive. The offspring was used to conduct immunohistochemistry experiments with brain slices. Data presented in Suppl. Fig. 10. This cross is considered “CR”. G, ACh neuron populations were genetically divided into a GABAergic subpopulation, Gad2-IRES-Cre mice were crossed with ChAT-IRES-FlpO and RC::FLTG Cre, FlpO reporter mice. Cells that expressed at one point ChAT were tdT positive. Cells that expressed at one point both ChAT and Gad65 were eGFP-positive. The offspring was used to conduct immunohistochemistry experiments with brain slices. Data presented in Fig. 7. This cross is considered “CGR”. H, ACh neuron populations were genetically divided into a GABAergic subpopulation, ChAT-IRES-Cre mice were crossed with vGAT-IRES2-FlpO-D and RC::FLTG Cre, FlpO reporter mice. FlpO-expressing cells were tdTomato-positive were td positive. FlpO- and Cra-expressing cells were eGFP-positive. The offspring was used to conduct immunohistochemistry experiments with brain slices. Data presented in Fig. 7. This cross is considered “CVR”.

**Suppl. Figure 2: Characterization of labeling efficiency of ACh neurons using the ChAT-FlpO x Ai65F cross.**

A, Animal model used to label cells positive for ChAT with tdT (green and magenta). B-J, 50 µm horizontal brain sections of mice expressing tdT in ChAT neurons were obtained at 2 months of age and counterstained for ChAT. Sections were imaged for tdT (green) and ChAT labelled with A647 (magenta). These brain regions are depicted: caudoputamen (CPu, B), oculomotor & trochlear nerve (CN III/IV, C), laterodorsal tegmentum and pedunculopontine nucleus (LDT/PPN, D), medial habenula (mHb, E), cerebral cortex (ctx, F), medial septum and diagonal band of Broca (MS/NDB, G), trigeminal nerve (CN V, H), facial nerve (CN VII, I), and vagus/hypoglossal nerve (CN X/XII, J). Insets depict magnified ROIs (small white square). Scale bar: 100 µm. K-M, Three-dimensional visualization of three cleared mouse brains hemispheres at 2 months of age with tdT-labelled ACh neurons for CAF. Scale bar: 1 mm.

**Suppl. Figure 3: Overview of all cleared WT and vGAT CKO mouse brains imaged with the light-sheet microscope.**

A-R, Three-dimensional visualization of all cleared vGAT WT and vGAT CKO mouse brains imaged with the light-sheet microscope at 2 months of age. Each brain is named by genotype (WT or CKO), litter number, animal number, and sex (m or f).

**Suppl. Figure 4: Overview of all cleared CGA mouse brains imaged with the light-sheet microscope.**

A-R, Three-dimensional visualization of all cleared CGA mouse brains imaged with the light-sheet microscope at 2 months of age. Each brain is named by litter number, animal number, and sex (m or f).

**Suppl. Figure 5: Overview of all cleared CVA mouse brains imaged with the light-sheet microscope.**

A-R, Three-dimensional visualization of all cleared CVA mouse brains imaged with the light-sheet microscope at 2 months of age. Each brain is named by litter number, animal number, and sex (m or f).

**Suppl. Figure 6: Cleared tissue count data quality control parameter verification in WT.**

A-E, Distribution of segmented cell counts by cell volume (pixels) at thresholds between 140K and 260K for WT in caudoputamen (A), isocortex (B), medial habenula (C), nucleus accumbens (D), and substantia innominata (E). Points within the gray rectangle were within defined minimal and maximal cell volume values and were used in cell volume analysis. F, Mean cell volumes of major brain regions by animal and genotype (open symbols). Estimated mean cell volumes for major brain regions and genotypes with 95% confidence intervals from a mixed model (filled symbols).

**Suppl. Figure 7: Differences in cell counts between ACh and ACh/GABA populations are robust independent of segmentation threshold.**

Mean cell counts by major brain region at each threshold for WT, CKO, and CGA (140K – 260K; WT n=9, CKO n=9, CGA m n=9, CGA f n=9 samples), as well as CVA (20K-260K for four samples, 20K-100K for the remaining 14 samples). Boxplots were color coded by WT (green), CKO (yellow), CGA (magenta), and CVA (turquoise). From left to right, top to bottom: cortical subplate, hypothalamus, isocortex, medulla, midbrain, olfactory areas, pallidum, pons, striatum, and thalamus. Boxes indicate interquartile range ($25^{th}$-$75^{th}$ percentiles); whiskers extend to minimum and maximum values (excluding outliers) and outliers as open circles.

**Suppl. Figure 8: Heatmap of all brain regions with more than 25 ACh neurons for all cleared WT, CKO, CGA, and CVA brains.**

Heatmap of cell counts of 54 brains (columns) by 108 composite regions (rows) with low cell counts in dark green and high cell counts in dark brown. Hierarchical clustering revealed that WT, CKO, and CGA brains form one cluster and CVA brains form a second cluster, and that the composite brain regions cluster into 4 distinct groups.

**Suppl. Figure 9: No sex differences for ACh neuron sub-populations that co-express GABAergic marker Gad2 or vGAT.**

Heatmaps of cell counts of individual (A) CGA brains and (C) CVA brains by sex (rows) and major brain region (columns), with low cell counts in dark green and high cell counts in dark brown. Volcano plots for (B) CGA and (D) CVA of statistical significance (expressed as -log10 FDR-adjusted p-value or log worth) by estimated multiplicative change in cell count for females relative to males (rate ratio) for 108 composite regions. Horizontal dashed line indicates significance threshold (FDR-adjusted p = 0.01); vertical dashed line indicates no change (rate ratio = 1).

**Suppl. Figure 10: Characterization of labeling efficiency of ACh neurons using the ChAT-FlpO x RC::FLTG cross.**

A, Animal model used to label cells positive for ChAT with tdT (green and magenta) while not labeling ChAT-negative neurons (grey). B-I, 50 µm horizontal brain sections of mice expressing tdT in ChAT/Gad2 neurons were obtained at 2 months of age and counterstained for ChAT. Sections were imaged for tdT (green) and ChAT labelled with A647 (magenta). These brain regions are depicted: caudoputamen (CPu, B), oculomotor & trochlear nerve (CN III/IV, C), laterodorsal tegmentum and pedunculopontine nucleus (LDT/PPN, D), medial septum and diagonal band of Broca (MS/NDB, E) nucleus basalis of Meynert and globus pallidus (MA/GP, F), cerebral cortex (ctx, G) trigeminal nerve (CN V, H), facial nerve (CN VII, I). Insets depict magnified ROIs (small white square). Scale bar: 100 µm.

### Suppl. Figure 1:

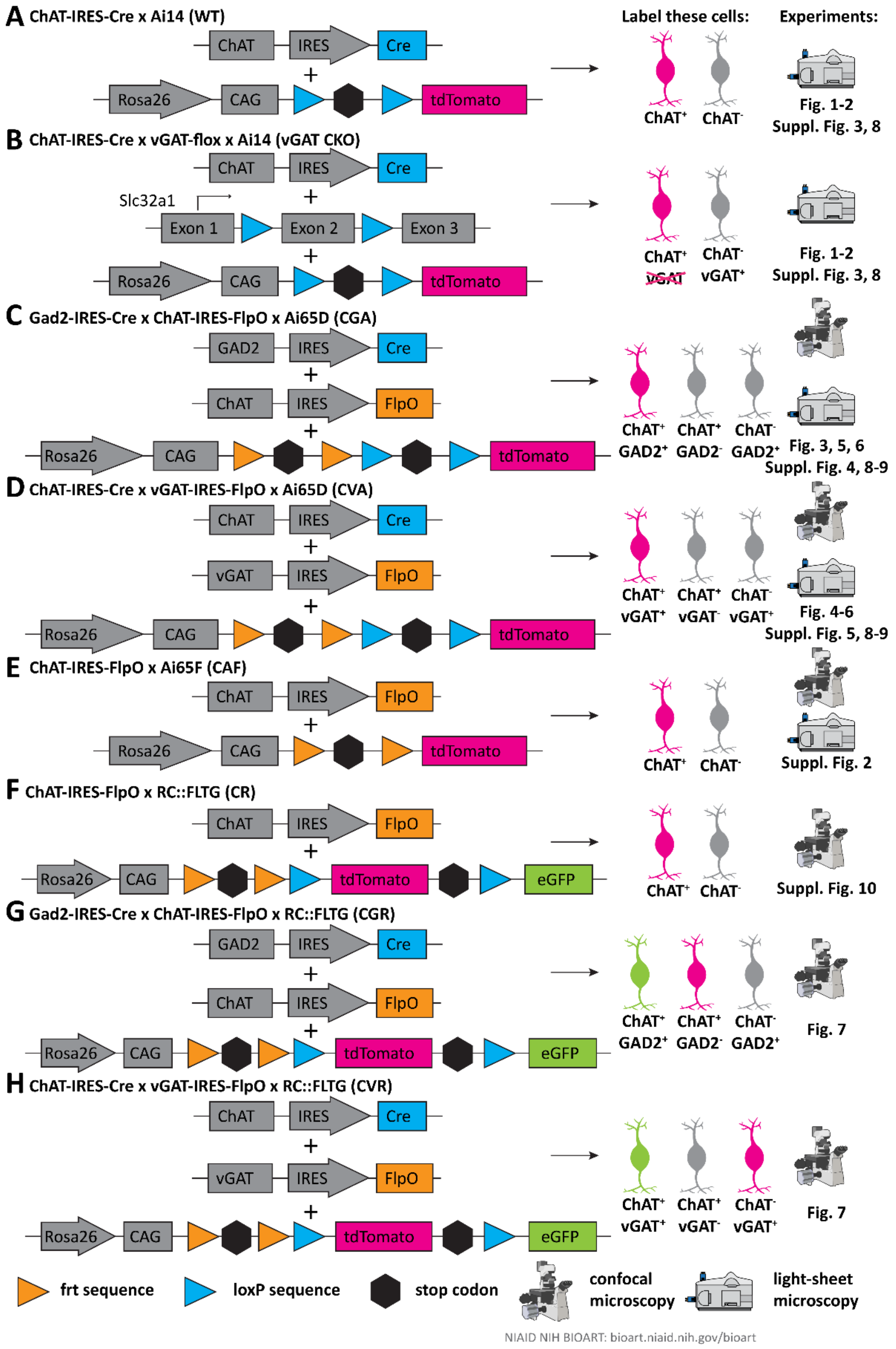

**Suppl. Figure 2:**

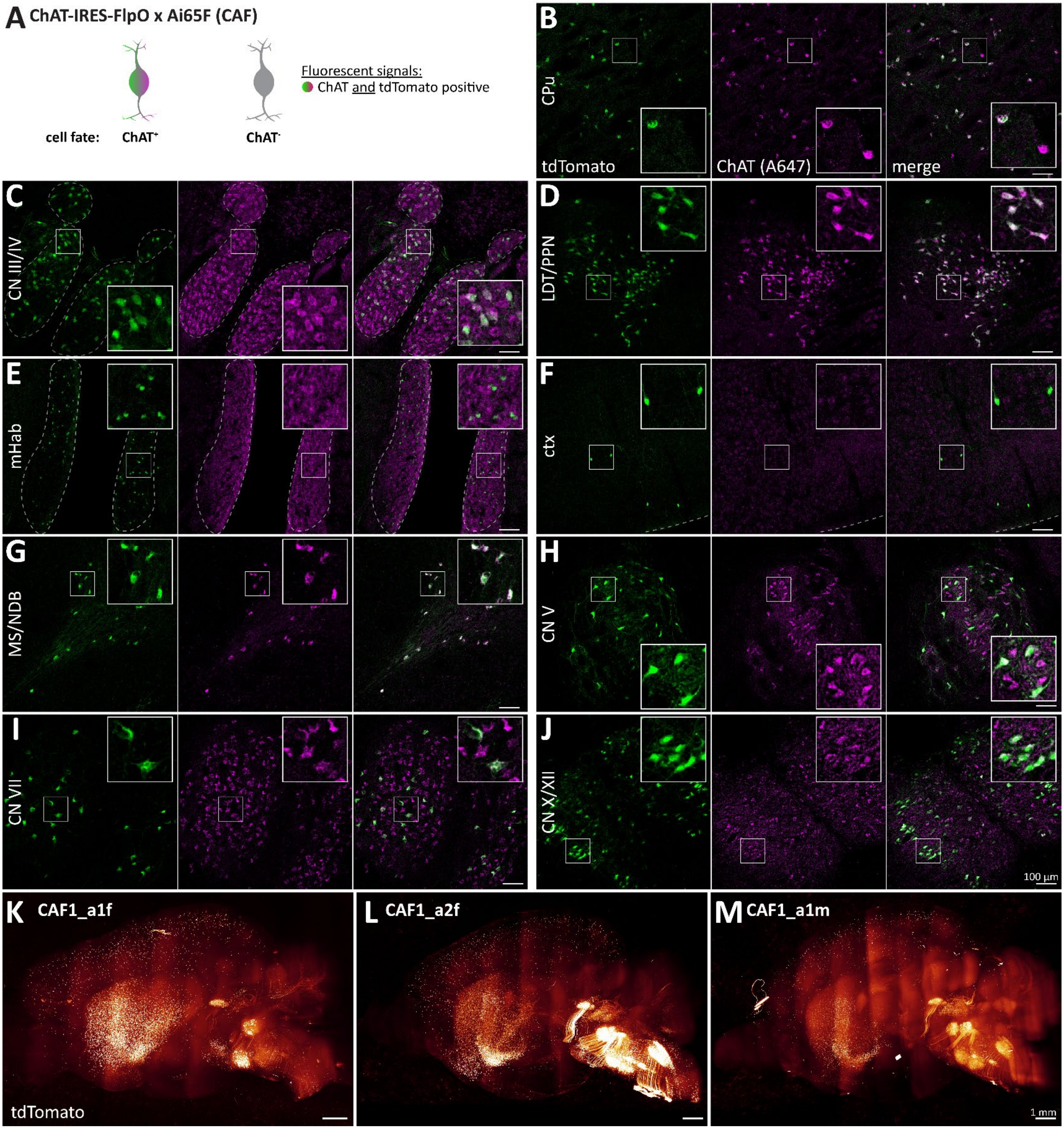

**Suppl. Figure 3:**

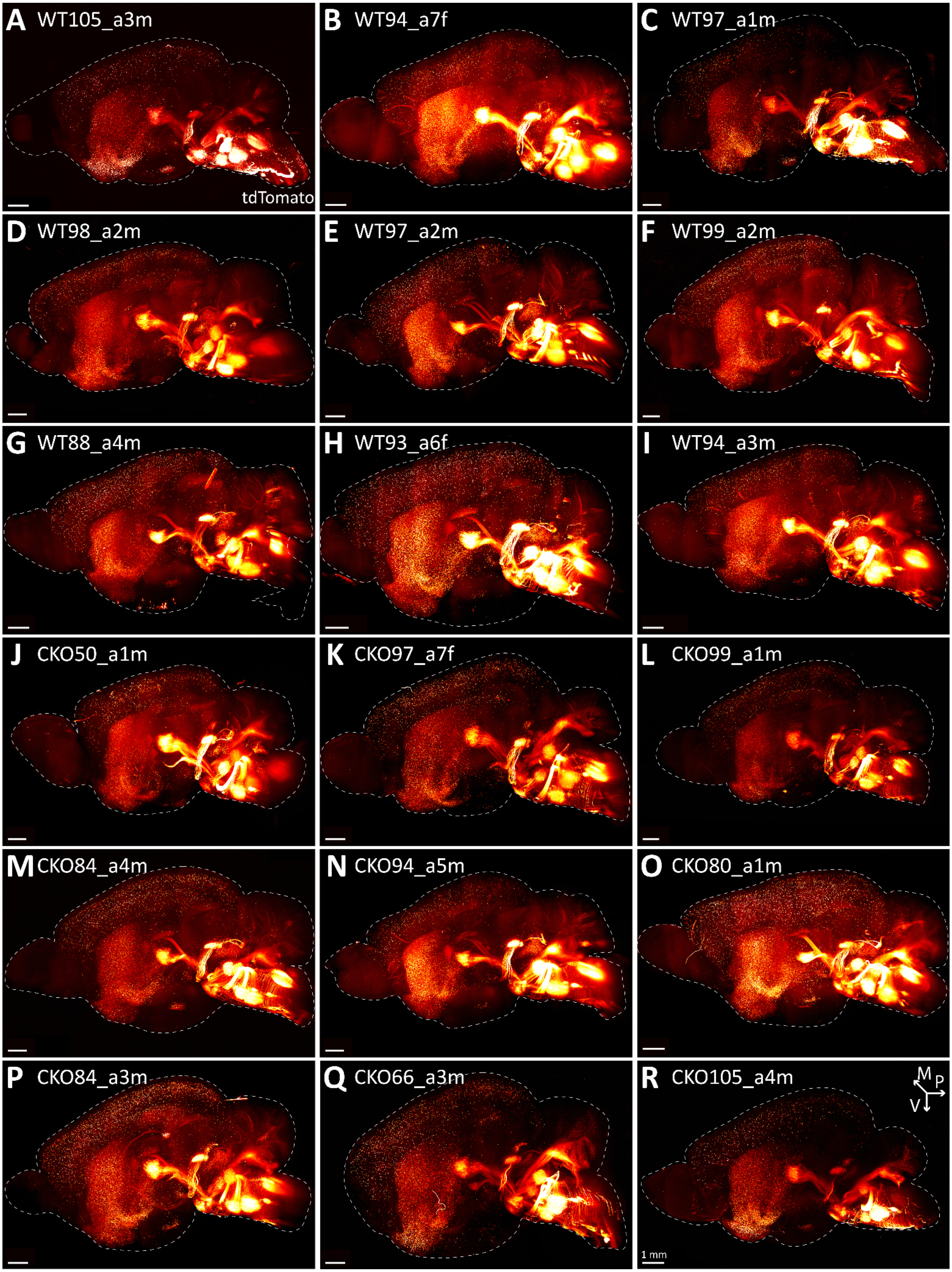

**Suppl. Figure 4:**

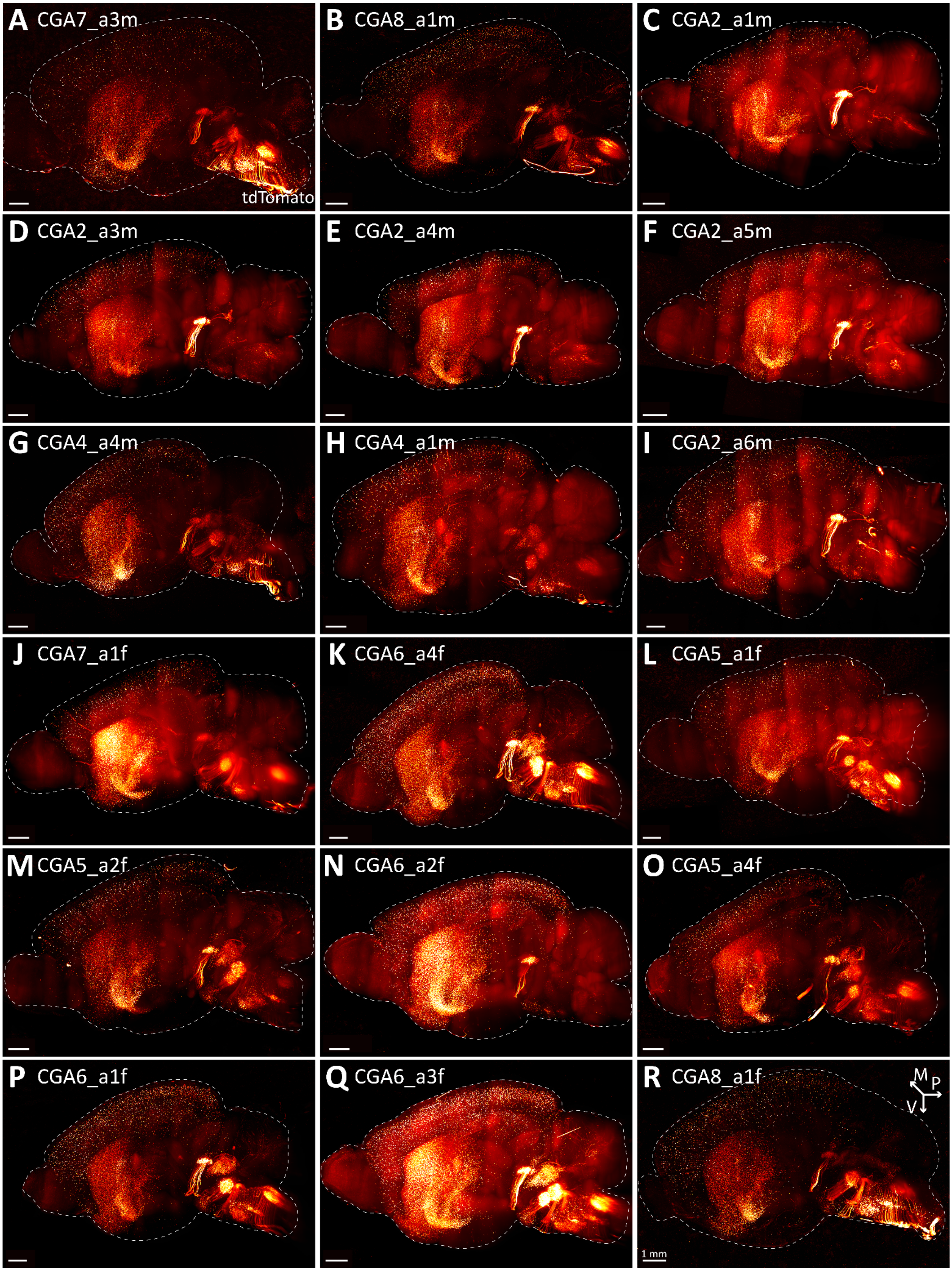

**Suppl. Figure 5:**

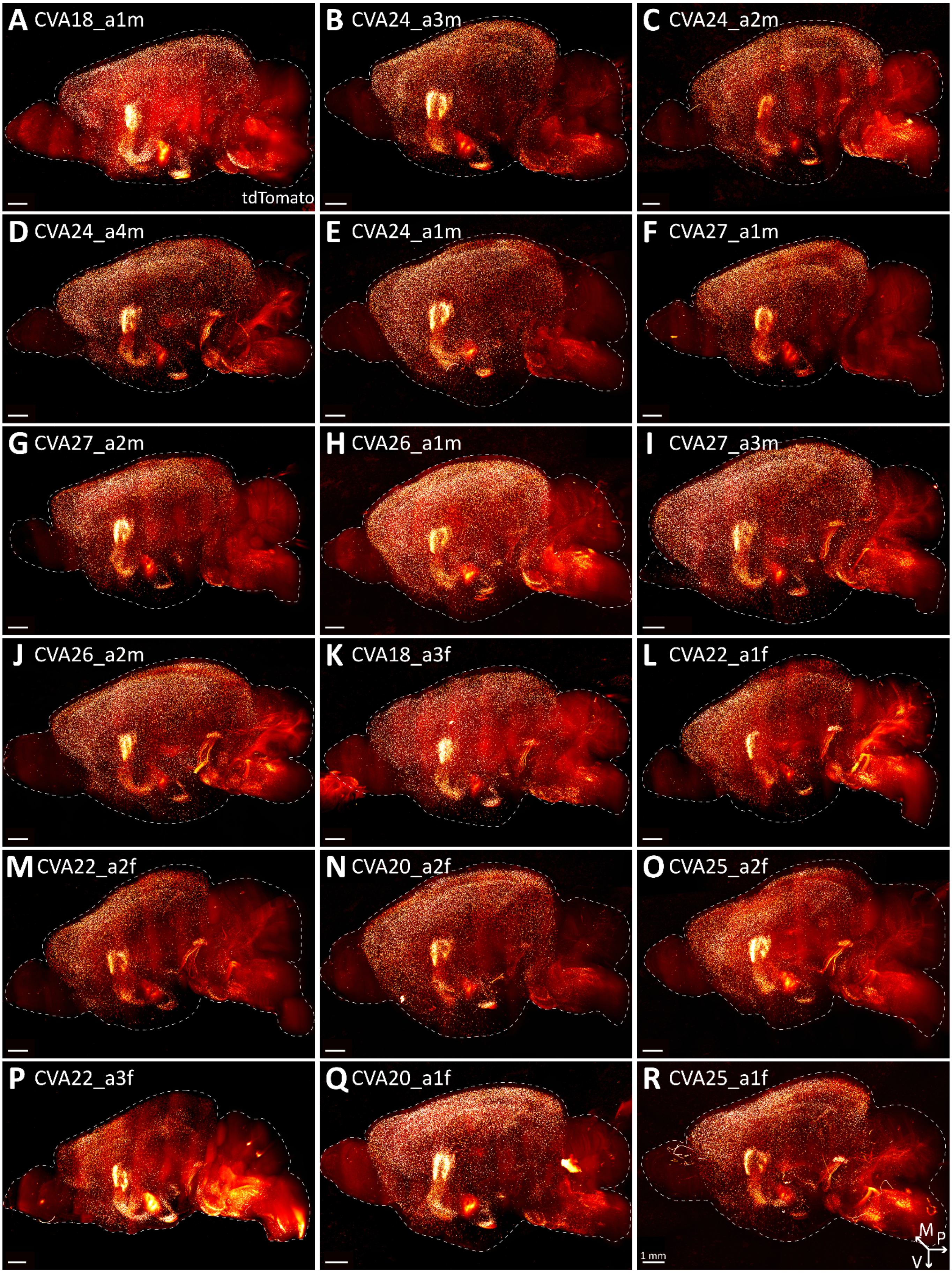

**Suppl. Figure 6:**

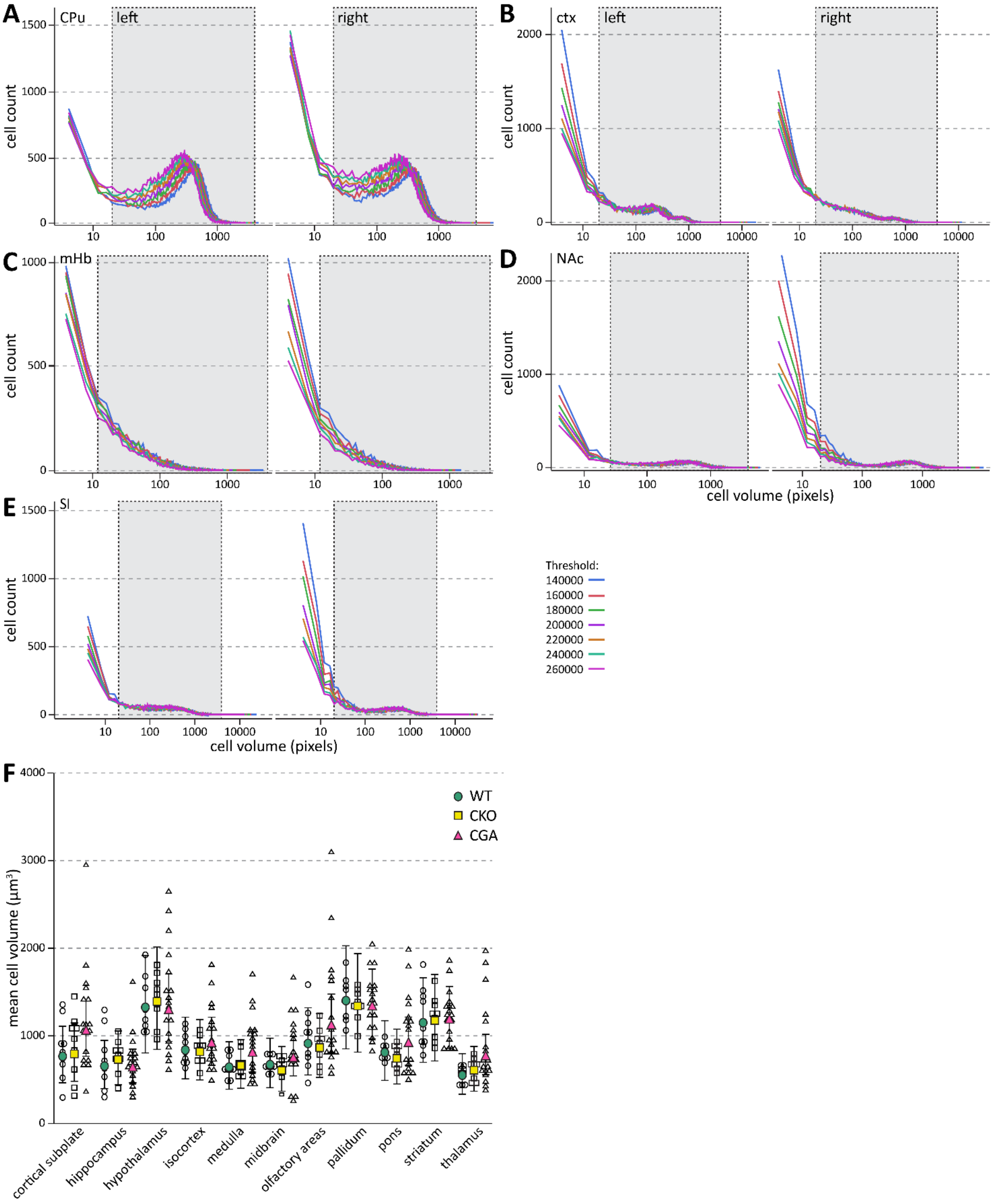

**Suppl. Figure 7:**

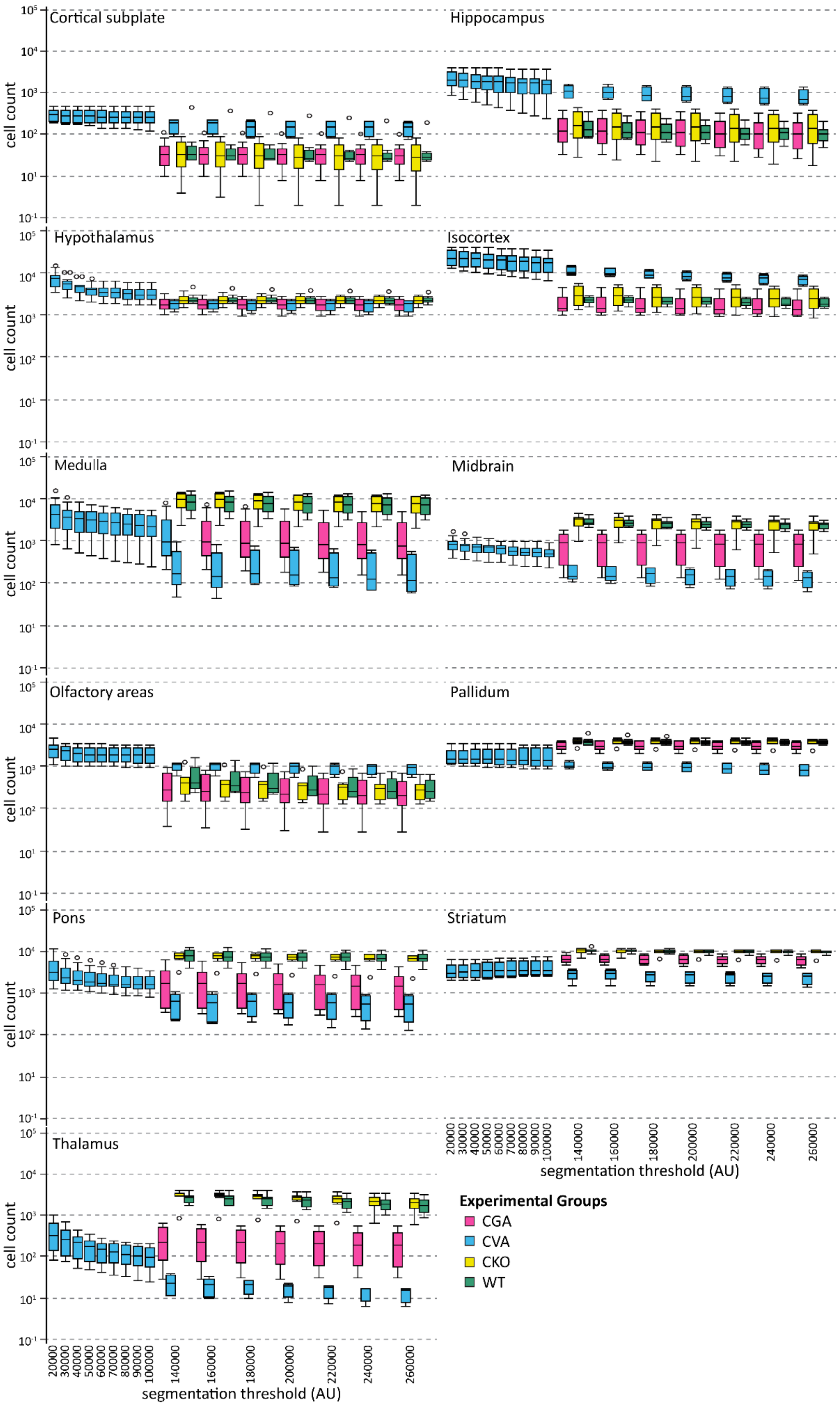

**Suppl. Fig. 8:**

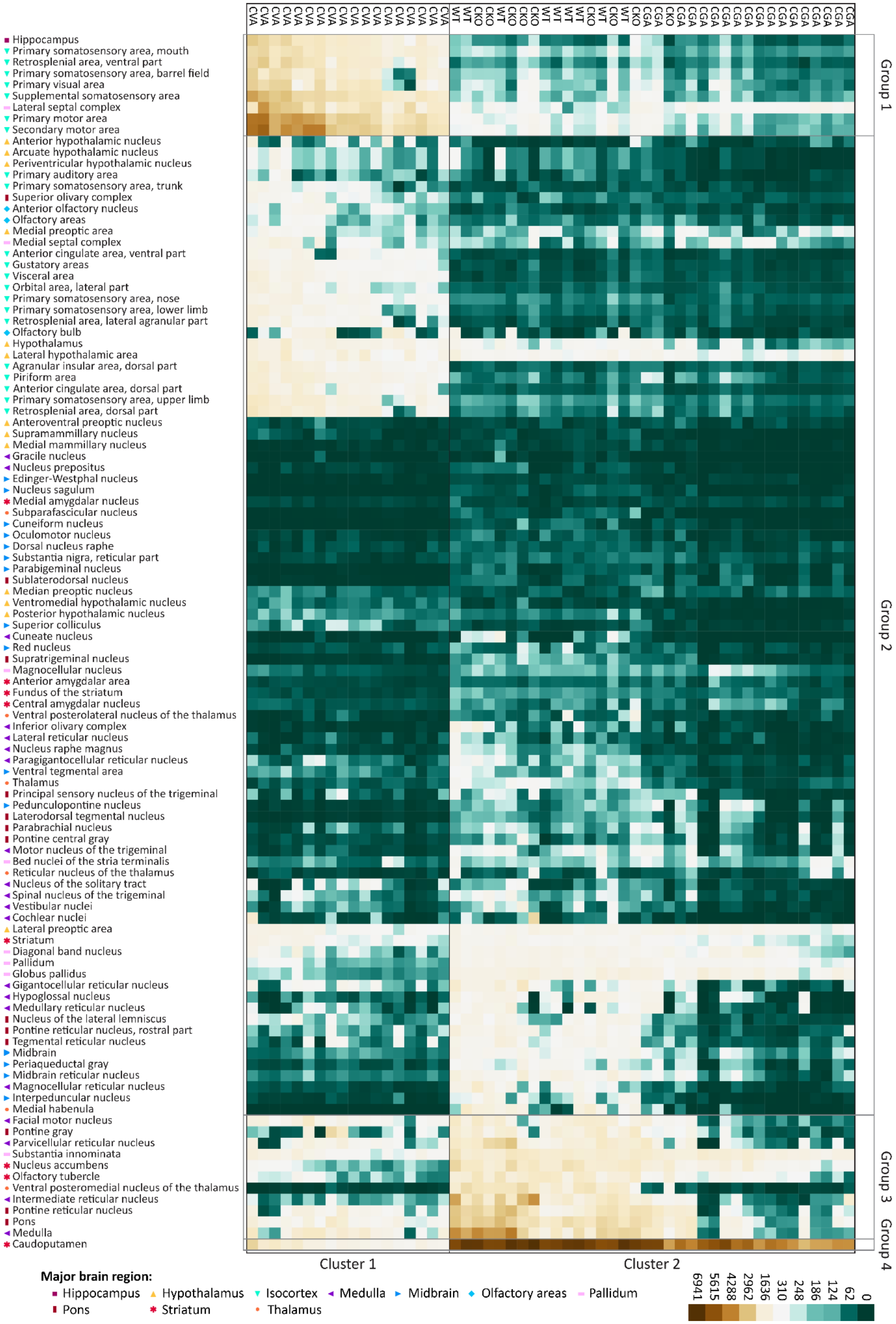

CVA CVA CVA CVA CVA CVA CVA CVA CVA CVA CVA CVA CVA CVA CVA CVA CVA CVA
WT WT CKO CKO WT CKO CKO CKO WT WT WT WT CKO WT CKO WT CKO CGA CGA CKO CGA CGA CGA CGA CGA CGA CGA CGA CGA CGA CGA CGA CGA CGA CGA CGA
Hippocampus
Primary somatosensory area, mouth
Retrosplenial area, ventral part
Primary somatosensory area, barrel field
Primary visual area
Supplemental somatosensory area
Lateral septal complex
Primary motor area
Secondary motor area
Anterior hypothalamic nucleus
Arcuate hypothalamic nucleus
Periventricular hypothalamic nucleus
Primary auditory area
Primary somatosensory area, trunk
Superior olivary complex
Anterior olfactory nucleus
Olfactory areas
Medial preoptic area
Medial septal complex
Anterior cingulate area, ventral part
Gustatory areas
Visceral area
Orbital area, lateral part
Primary somatosensory area, nose
Primary somatosensory area, lower limb
Retrosplenial area, lateral agranular part
Olfactory bulb
Hypothalamus
Lateral hypothalamic area
Agranular insular area, dorsal part
Piriform area
Anterior cingulate area, dorsal part
Primary somatosensory area, upper limb
Retrosplenial area, dorsal part
Anteroventral preoptic nucleus
Supramammillary nucleus
Medial mammillary nucleus
Gracile nucleus
Nucleus prepositus
Edinger-Westphal nucleus
Nucleus sagulum
Medial amygdalar nucleus
Subparafascicular nucleus
Cuneiform nucleus
Oculomotor nucleus
Dorsal nucleus raphe
Substantia nigra, reticular part
Parabigeminal nucleus
Sublaterodorsal nucleus
Median preoptic nucleus
Ventromedial hypothalamic nucleus
Posterior hypothalamic nucleus
Superior colliculus
Cuneate nucleus
Red nucleus
Supratrigeminal nucleus
Magnocellular nucleus
Anterior amygdalar area
Fundus of the striatum
Central amygdalar nucleus
Ventral posterolateral nucleus of the thalamus
Inferior olivary complex
Lateral reticular nucleus
Nucleus raphe magnus
Paragigantocellular reticular nucleus
Ventral tegmental area
Thalamus
Principal sensory nucleus of the trigeminal
Pedunculopontine nucleus
Laterodorsal tegmental nucleus
Parabrachial nucleus
Pontine central gray
Motor nucleus of the trigeminal
Bed nuclei of the stria terminalis
Reticular nucleus of the thalamus
Nucleus of the solitary tract
Spinal nucleus of the trigeminal
Vestibular nuclei
Cochlear nuclei
Lateral preoptic area
Striatum
Diagonal band nucleus
Pallidum
Globus pallidus
Gigantocellular reticular nucleus
Hypoglossal nucleus
Medullary reticular nucleus
Nucleus of the lateral lemniscus
Pontine reticular nucleus, rostral part
Tegmental reticular nucleus
Midbrain
Periaqueductal gray
Midbrain reticular nucleus
Magnocellular reticular nucleus
Interpeduncular nucleus
Medial habenula
Facial motor nucleus
Pontine gray
Parvicellular reticular nucleus
Substantia innominata
Nucleus accumbens
Olfactory tubercle
Ventral posteromedial nucleus of the thalamus
Intermediate reticular nucleus
Pontine reticular nucleus
Pons
Medulla
Caudoputamen
Group 1
Group 2
Group 3
Group 4
Cluster 1
Cluster 2
Major brain region:
Hippocampus
Hypothalamus
Isocortex
Medulla
Midbrain
Olfactory areas
Pallidum
Pons
Striatum
Thalamus
6941
5615
4288
2962
1636
310
248
186
124
62
0

**Suppl. Figure 9:**

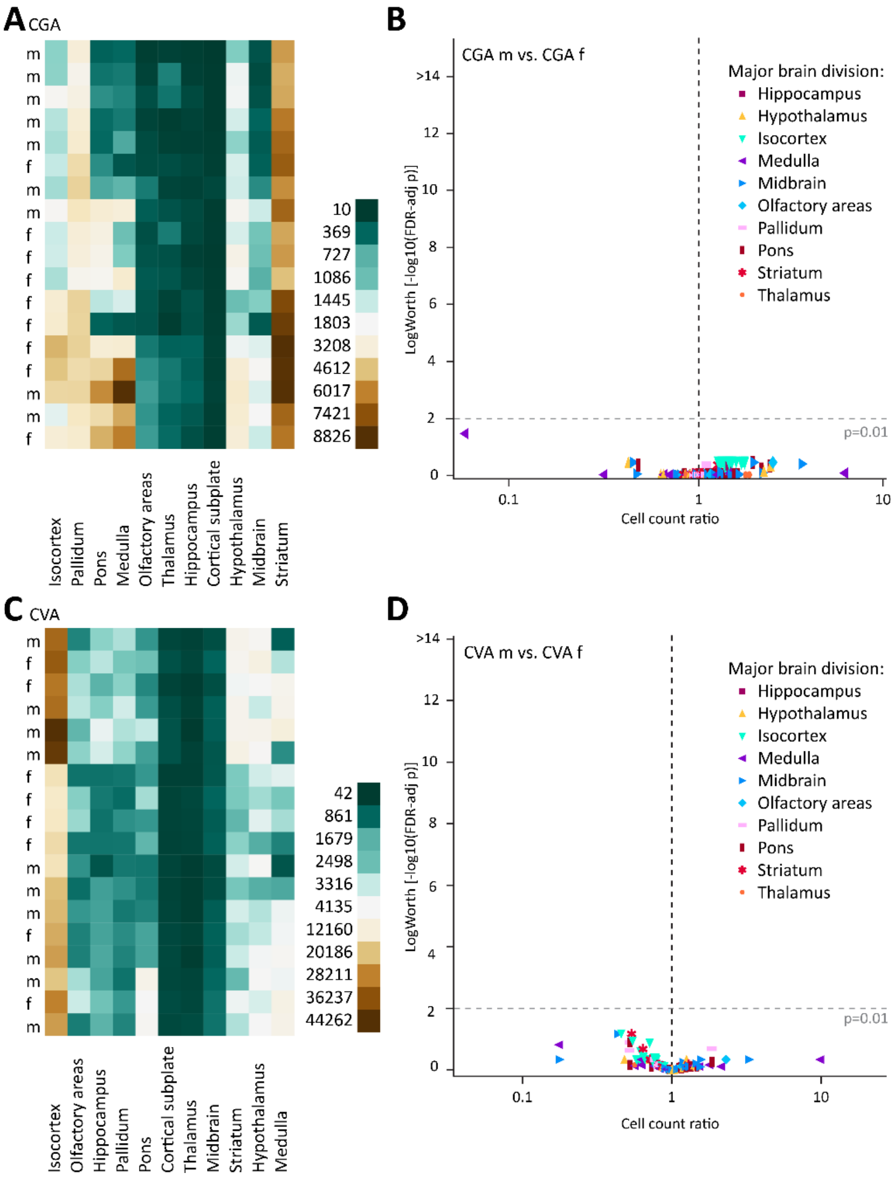

**Suppl. Figure 10:**

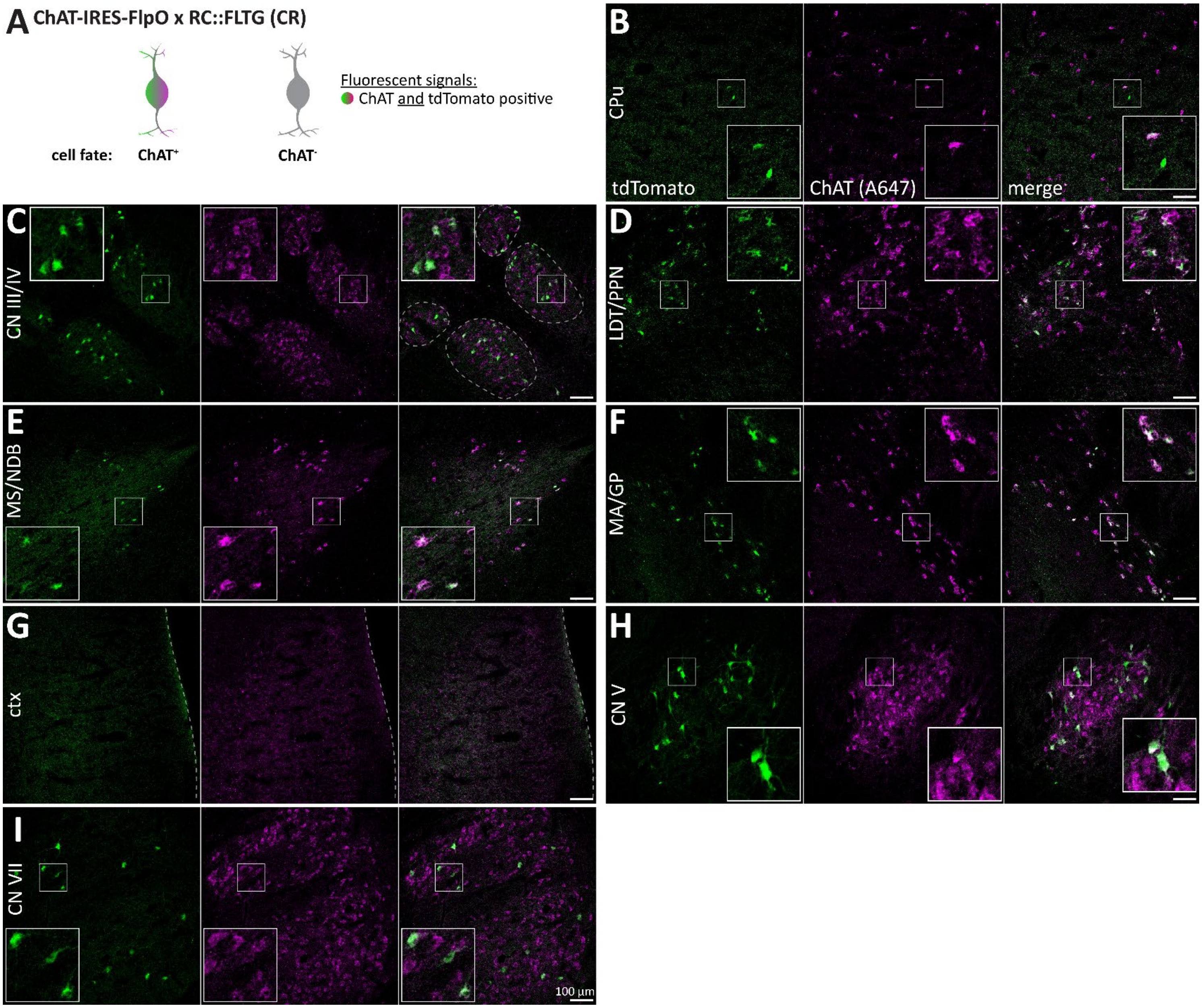
